# Silicon modulates multi-layered defense against powdery mildew in Arabidopsis

**DOI:** 10.1101/2020.02.05.935734

**Authors:** Lili Wang, Min Dong, Qiong Zhang, Ying Wu, Liang Hu, James F Parson, Edward Eisenstein, Xiangge Du, Shunyuan Xiao

## Abstract

Silicon (Si) has been widely employed in agriculture to enhance resistance against pathogens in many crop plants. However, the underlying molecular mechanisms of Si-mediated resistance remain elusive. In this study, the Arabidopsis-powdery mildew pathosystem was employed to investigate possible defense mechanisms engaged for Si-mediated resistance. Because Arabidopsis lacks efficient Si transporters and thus is a low Si-accumulator, two heterologous Si influx transporters (from barley and muskmelon) were individually expressed in wild-type Arabidopsis Col-0 and a panel of mutants defective in different immune signaling pathways. Results from infection tests showed that while very low leaf Si content slightly induced salicylic acid (SA)-dependent resistance, high Si promoted PAD4-dependent but EDS1- and SA-independent resistance against the adapted powdery mildew isolate *Golovinomyces cichoracearum* UCSC1. Intriguingly, our results also showed that high Si could largely reboot non-host resistance in an immune-compromised *eds1/pad4/sid2* triple mutant background against a non-adapted powdery mildew isolate *G. cichoracearum* UMSG1. Taken together, our results suggest that assimilated Si modulates distinct, multi-layered defense mechanisms to enhance plant resistance against adapted and no-adapted powdery mildew pathogens, possibly via synergistic interaction with defense-induced callose.

## INTRODUCTION

Silicon (Si) is the second most abundant element on the earth. Although it is not considered to be an essential element for plant growth, Si has long been recognized as a “beneficial" or “quasi-essential” substance to plants, mainly due to its important role in plant nutrition, particularly under stressful conditions. Over the past 25 years, a plethora of studies with >800 publications have collectively shown that Si can effectively protect plants from abiotic stresses, including drought, salinity and metals toxicity, as well as biotic stresses caused by insect herbivores or various pathogens ranging from viruses, bacteria, fungi, oomycetes, to nematodes (Van Bockhaven et al. 2012; Coskun et al. 2019). However, despite the extensive studies, the exact molecular mechanisms underlying the protective roles of Si in plants still remain elusive (Coskun et al. 2019). Conceivably, this knowledge gap limits the exploitation of the full potential concerning practical application of Si in agriculture.

It has been demonstrated that Si needs to be absorbed by plants through passive channel-type, selective Si transporters to realize its prophylactic effect (Ma 2010). Intriguingly, Si accumulation varies over 100-fold in different plant species, ranging from 0.1% to 10% in dry weight (Epstein 1994). Based on the levels of Si accumulation, plants are divided into three categories: high Si (or active) accumulator, intermediate Si (or passive) accumulator and low Si (rejective) accumulator (Takahashi et al. 1990). Understandably, most studies on Si transport and its physiological roles in plants have been conducted with high accumulators, and to a lesser extent, intermediate accumulators. The ability of a particular plant for accumulation of Si is determined by the transport efficiency of its Si influx transporters. The first identified Si influx transporter (*OsLsi1*) in higher plants was found in rice in 2006 (Ma et al. 2006). Since then, many Si transporters homologous to *OsLsi1* have been identified and characterized in different plant species, such as barley *(HvLsi1*) (Chiba et al. 2009), wheat (*TaLsi1*) (Montpetit et al. 2012), pumpkin (*CmLsi1*) (Mitani et al. 2011). By contrast, no similar Si transporter has been identified in Arabidopsis, agreeing with its classification as a low Si accumulator.

Mechanistic studies on how Si exerts prophylactic effect on plants using high or intermediate Si accumulators have been focused on the aspect of disease resistance. Based on published data generated with distinct pathosystems, two types of mechanisms have been proposed to explain how Si might act in plant cells. The first type conforms to the “apoplastic obstruction hypothesis” recently proposed by Coskun and colleagues based on many observations that Si is deposited on plant cell surface or in the apoplastic space whereby it strengthens physical barriers that can limit entry of pathogens and/or delivery of pathogen molecules (Coskun et al. 2019). For example, Si can accumulate and deposit beneath the cuticle of rice leaves to reduce penetration of rice blast *Magnaporthe grisea* (Yoshida 1965) and Si can also polymerize in specialized cells and cellular structures of some species (particularly grasses), such as leaf silica and long cells, and spikelet hairs and papillae to strengthen the physical barriers to prevent pathogen entry (Rafi et al. 1997). For the second type of mechanism, some studies have suggested that Si may be able to potentiate a plant defense program(s), which enables more sensitive and rapid response of plants to pathogen attack (Van Bockhaven et al. 2012). Inducible plant defense mechanisms have been extensively studied in the field of molecular plant-microbe interaction and known to engage a conserved and complex network of signal transduction pathways that may involve activation of mitogen-activated protein Kinases, and/or production and signaling of small-molecule phytohormones such as salicylic acid (SA), jasmonic acid (JA), and/or ethylene (ET) (Pieterse et al. 2009). If these molecular and biochemical events are “primed” to a more ready-state by Si, more effective chemical defense ensues. For example, Si application was reported to reprogram gene transcription in uninoculated rice plants (Van Bockhaven et al. 2012) and may modulate the rice ET pathway to induce resistance to the brown spot fungus *Cochliobolus miyabeanus*, which is a necrotrophic pathogen (Van Bockhaven et al. 2015). However, in many cases, Si supplementation had little, if any, effect on gene expression of plants (Fauteux et al. 2006; Brunings et al. 2009; Chain et al. 2009).

The role of Si in promoting disease resistance is particularly evident in the case of powdery mildew as shown in cucumber, muskmelon, and zucchini squash, barley, and rose (Menzies et al. 1992; Wiese et al. 2005; Shetty et al. 2012). In mechanistic studies concerning Si-mediated resistance to PM, Si has been shown to strengthen papilla-mediated inhibition of appressorial penetration in wheat (Bélanger et al. 2003), and it has also been reported to promote accumulation of antifungal flavonoid phytoalexins in rose (Shetty et al. 2011). Thus, Si appears to promote both physical and chemical barriers against powdery mildew (and possibly other) pathogens. However, the genetic and molecular basis of Si’s action largely remained unexplored until a recent genetic study in which transgenic Arabidopsis expressing a heterologous Si transporter was exploited to address this question. Vivancos and colleagues (2015) expressed the wheat Si transporter (*TaLsi1*) in Arabidopsis wild-type Col-0 and mutants that are defective in SA-signaling (due to the loss of PAD4) or biosynthesis (due to the loss of SID2). They showed that *TaLsi1*-expressing Col-0 plants were able to accumulate high levels of Si and hence exhibited enhanced resistance to powdery mildew. Interestingly, they found that *pad4* and *sid2* mutant plants transgenic for *TaLsi1* also showed similar levels of resistance when Si was supplied. These results suggested that Si, when above a threshold level, is able to activate disease resistance via an unknown mechanism(s) that is independent of SA signaling (Vivancos et al. 2015).

In this study, we employed the Arabidopsis-powdery mildew pathosystem to further investigate the molecular mechanisms of Si-mediated disease resistance in plants. Through the use of Arabidopsis genetic mutants, transgene expression of heterologous Si transporters and a more stringent control of Si supply, we found that Arabidopsis plants containing either very low or very high Si content displayed enhanced resistance to powdery mildew. Interestingly, we found that whereas very low Si-induced resistance is SA-dependent, high Si-mediated resistance is indeed independent of SA signaling but surprisingly requires PAD4. Moreover, we also found that high Si can effectively reboot penetration resistance in a SA signaling-defective and *PAD4*-ablated Arabidopsis mutant that is otherwise fully susceptible to a non-adapted powdery mildew isolate. Thus, our results present genetic evidence for the role of Si in multi-layered plant defense mechanisms, which should help clarify the current confusion and controversy over the molecular mechanisms underlying Si’s positive role in plant disease resistance against fungal pathogens.

## RESULTS

### Plants with either very low or very high endogenous Si display enhanced resistance against powdery mildew

In our pilot experiment, we surprisingly found that *Arabidopsis thaliana* wild-type Col-0 plants became slightly more susceptible to an adapted powdery mildew isolate *Golovinomyces cichoracearum* (*Gc*) UCSC1 when irrigated water containing 1.7 mM Silicon. This is an equivalent concentration of Si that provided remarkable protection against *B. graminis f. sp. tritici* in wheat (Chain et al. 2009) compared to plants without Si (Figure S1). To reliably assess the role of Si in modulating plant defense mechanisms, we decided to use perlite irrigated with deionized pure water containing Si and a controlled amount of fertilizer to grow Arabidopsis plants (Xiao et al. 2003). We first tested eight-week-old wild-type Col-0 plants with *Gc* UCSC1. Intriguingly, we found that Col-0 plants irrigated with nutrient solution without Si (–Si) showed enhanced disease resistance, which was visible by the naked eye when compared with plants irrigated with water and fertilizer plus 1.0 mM Si (+Si) (Figure 1A). Quantification of fungal spore production at 10 days post-inoculation (dpi) showed that plants of Col-0/–Si produced ~30% less spores than plants of Col-0/+Si (Figure 1B). Leaf Si content in Col-0/–Si plants was 1.60 mg g^−1^ dry leaf tissue, whereas Col-0/+Si plants had slightly but significantly higher leaf Si content (2.38 mg g^−1^; ~48% increase) (Figure 1C). These results reinforce the notion that Arabidopsis was a low Si-accumulator with poor Si absorption (Vivancos et al. 2015) and suggest that very low endogenous Si content may activate plant defense against powdery mildew.

**Figure 1.**
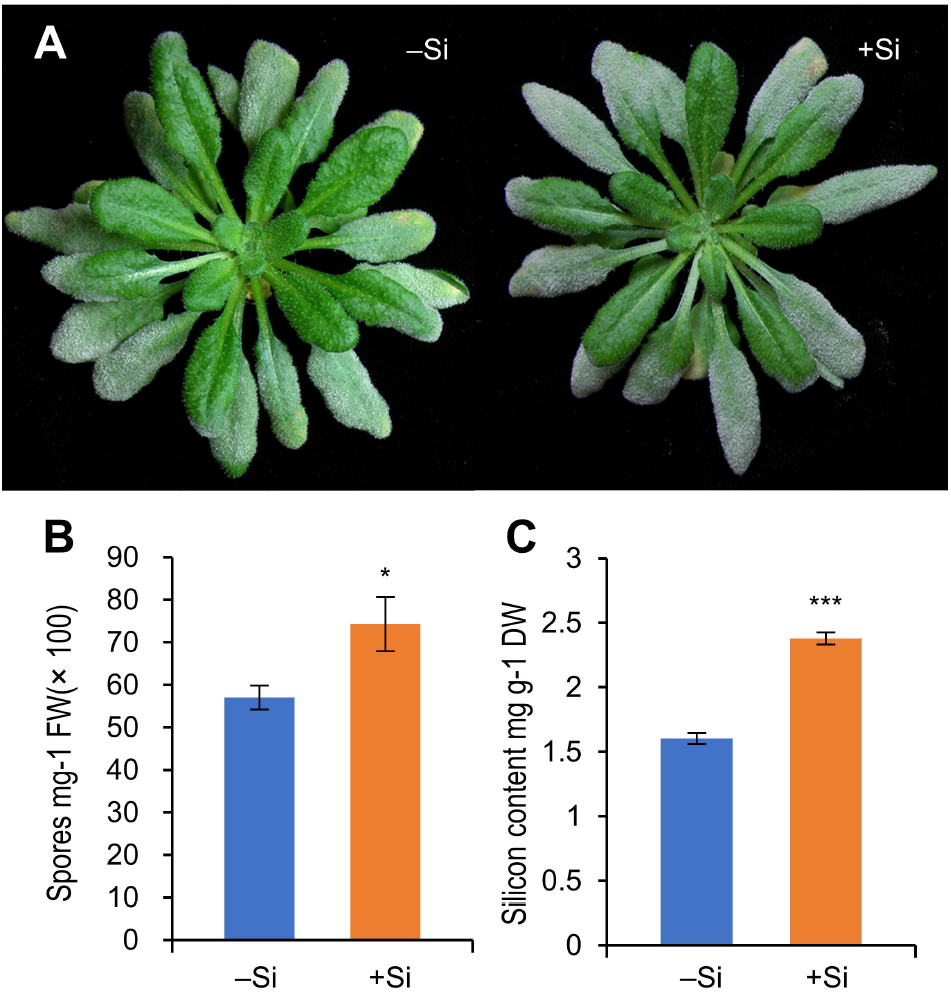
Low level of endogenous Silicon (Si) reduces resistance against well-adapted powdery mildew *Golovinomyces cichoracearum* (*Gc*) UCSC1 in *Arabidopsis thaliana* Col-0. (A) Representative images of Arabidopsis Col-0 infected with *Gc* UCSC1 at 10 dpi with treatment either 0 mM (–Si) or 1.0 mM (+Si) Si. (B) Quantification of spore production in the indicated treatments at 10 dpi normalized to leaf fresh weight. (C) Silicon content in leaves of Arabidopsis Col-0 treated with either 0 mM (–Si) or 1.0 mM (+Si) Si normalized to leaf dry weight. Bars represent standard errors and within treatments, and asterisks on a column denote a significant difference (*t*-test, *P<0.05, ***P<0.001).

In order to evaluate the impact of higher levels of assimilated Si in plant defense using Arabidopsis, we generated Col-0 lines expressing the barley (*Hordeum vulgare*) Si influx transporter-encoding gene *HvLsi1* (Chiba et al. 2009) or the muskmelon (*Cucumis melo*) Si influx transporter-encoding gene *CmeLsi1* (which was amplified from the genomic DNA of muskmelon based on sequence homology to the *Cucurbita moschata* Si influx transporter gene *CmLsi1*) (Mitani et al. 2011). As expected, all transgenic lines supplemented with Si showed high Si content. For example, *CmeLsi1*-expressing Col-0 plants of a representative T3 homozygous line treated with exogenous Si increased leaf Si content by ~280% (5.52±0.43 mg g^−1^) compared with untreated control plants of the same transgenic line (Figure 2A). Interestingly, HvLsi1, when expressed in Col-0, appeared to have a stronger Si absorbing capability than CmeLsi1, because leaf Si content in plants of one representative Col-0 transgenic line expressing *HvLsi1* increased by ~55% compared with untreated Col-0 even without exogenous Si application and further increased by ~318% (10.38±0.22 mg g ^−1^) in plants supplied with Si, resulting in Si accumulation of more than 1% of the total dry weight (Figure 2A). Similar patterns were seen in other genetic backgrounds (Figure S2).

**Figure 2.**
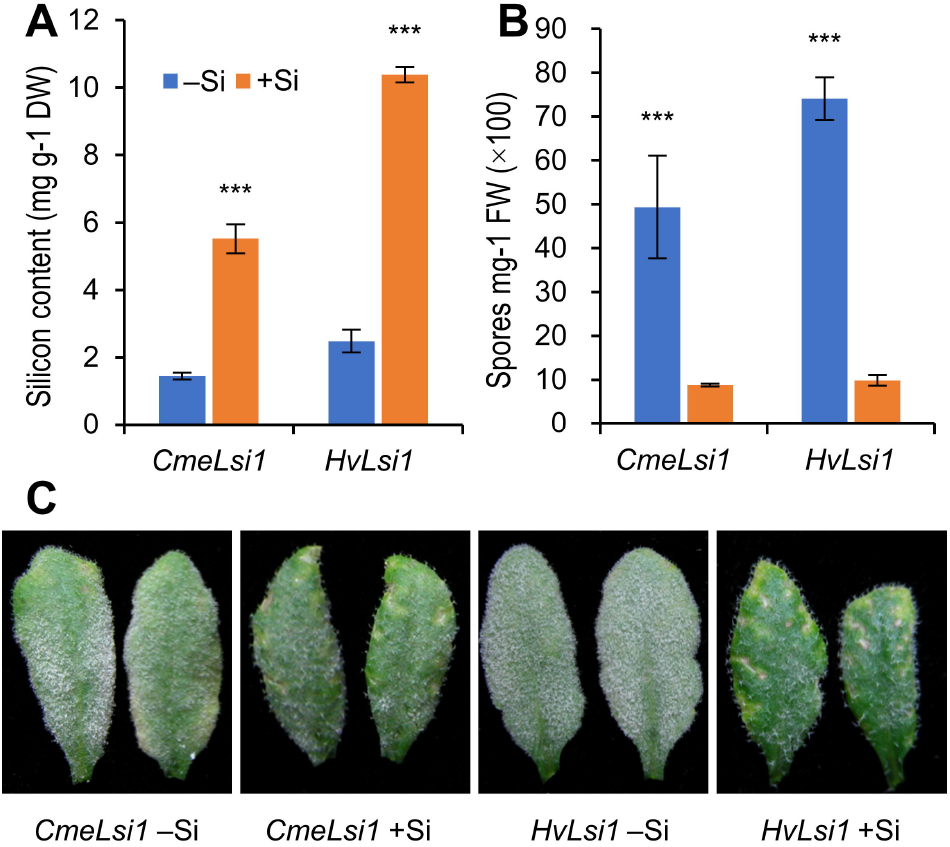
High level of endogenous Silicon (Si) enhances resistance against well-adapted powdery mildew *Golovinomyces cichoracearum* (*Gc*) UCSC1 in Col-0 overexpressing *CmeLsi1* or *HvLsi1*. (A) Silicon content in leaves of Arabidopsis *CmLsi1*/Col-0 or *HvLsi1*/Col-0 treated with either 0 mM (–Si) or 1.0 mM (+Si) Si normalized to leaf dry weight. (B) Quantification of spore production in the indicated treatments at 10 dpi normalized to leaf fresh weight. (C) Representative images of transgenic Arabidopsis Col-0 infected with *Gc* UCSC1 at 10 dpi with treatment either –Si or +Si. Bars represent standard errors and within treatments, and asterisks on a column denote a significant difference (*t*-test, ***P<0.001).

We then tested plants of these transgenic lines grown in perlite supplied with Si (1.0 mM) or without Si with *Gc* UCSC1. Visual disease scoring and spore quantification showed that both *HvLsi1*- and *CmeLsi1*-transgenic plants with Si supplement displayed remarkable enhanced resistance in comparison with plants of the respective transgenic lines without Si supplement (Figure 2C). Interestingly, *HvLsi1*-Col-0/–Si plants (with a leaf Si content of 2.49±0.34 mg g^−1^) showed a similar level of disease susceptibility as Col-0/+Si plants (with a similar leaf Si content of 2.38± 0.05 mg g ^−1^) (Figures 1A, 2C), whereas *CmeLsi1*-Col-0/–Si plants (with a leaf Si content of 1.45±0.10 mg g ^−1^) showed reduced susceptibility similar to the level seen in Col-0/–Si plants (with a leaf Si content of 1.60±0.04 mg g ^−1^) (Figures 1A, 2C). Quantification of fungal spore production fully supported the visual phenotypes (Figure 2B).

### Very low Si-conditioned resistance requires SA-pathway components EDS1, PAD4 and SID2

H_2_O_2_ production and accumulati**on** in mildew-invaded epidermal cells correlates with known powdery mildew resistance mechanisms in Arabidopsis (Xiao et al. 2003). To understand why plants with very low Si content are more resistant to PM, we first performed DAB (3,3’-Diaminobenzidine) staining of infected leaves of Col-0 for *in situ* detection of H_2_O_2_ accumulation. We found that Col-0/–Si plants showed more frequent H_2_O_2_ accumulation in PM-invaded epidermal cells (~15%) in comparison with Col-0/+Si plants (~5%) (Figure 3A), which likely contributed to the enhanced resistance in Col-0/–Si plants.

**Figure 3.**
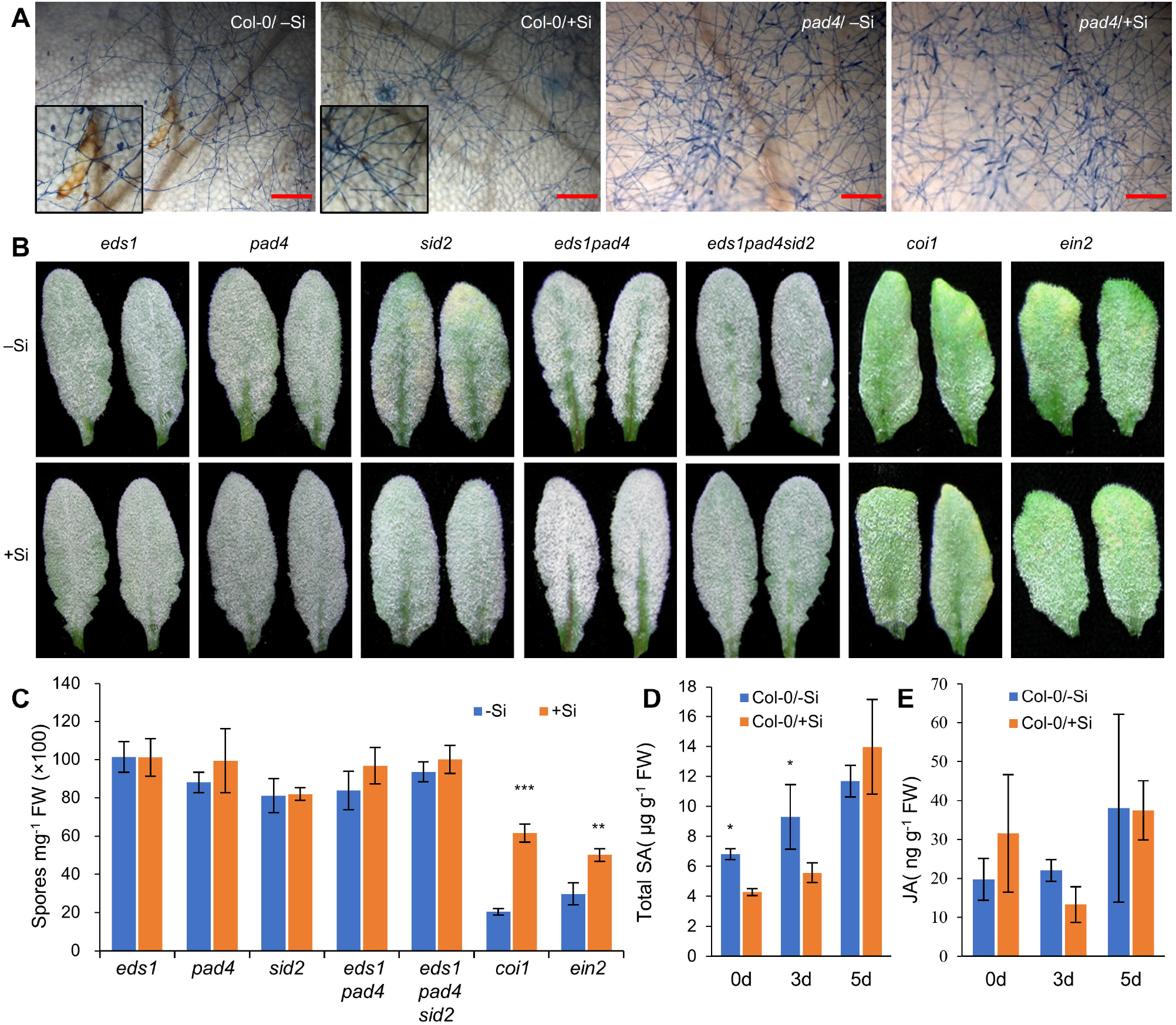
Very low Silicon(Si)-conditioned resistance requires SA-pathway components EDS1, PAD4 and SID2. (A) Representative microscopic images of typical *Gc* UCSC1 fungal microcolonies grown on leaves of the indicated genotypes with treatment either 0 mM (–Si) or 1.0 mM (+Si) Si at 5 dpi. H_2_O_2_ reacts with 3,3-diaminobenzidine to form a reddish-brown stain, while fungal structures are stained blue by trypan blue. Scale bars=200 μm. (B) Representative images of Arabidopsis leaves of the indicated genotypes infected with *Golovinomyces cichoracearum* (*Gc*) UCSC1 with treatment either 0 mM (–Si) or 1.0 mM (+Si) Si at 10 dpi. (C) Quantification of spore production in the indicated genotypes with treatment either 0 mM (–Si) or 1.0 mM (+Si) Si at 10 dpi normalized to leaf fresh weight. (D, E) Total SA and JA levels in leaves of wild-type Arabidopsis Col-0 with treatment either 0 mM (–Si) or 1.0 mM (+Si) Si at 0, 3 and 5 dpi. Bars represent standard errors and within treatments, and asterisks on a column denote a significant difference (t-test, *P<0.05, **P<0.01, ***P<0.001).

Next, to investigate the possible defense signaling pathway(s) engaged for the H_2_O_2_ production and resistance to powdery mildew due to very low Si content in Col-0/–Si plants, we prepared plants of *eds1*, *pad4*, *sid2*, *coi1*, *ein2* single and *eds1pad4* double, and *eds1pad4sid2* triple Arabidopsis mutants that are defective in SA-dependent (EDS1, PAD4 and SID2), jasmonic acid (JA)-dependent (COI1) and ethylene-dependent (EIN2) signaling pathways in perlite supplemented with or without Si. Infection tests with *Gc* UCSC1 revealed that plants of *coi1*/–Si and *ein2*/–Si showed slightly enhanced resistance in comparison with their respective +Si plants, similar to the situation of Col-0/–Si vs. Col-0/+Si (Figure 3B). By contrast, plants of the remaining genotypes that are defective in SA signaling or biosynthesis displayed no significant difference between –Si and +Si-treated plants (Figure 3B). Quantification of fungal spore production supported the visual phenotypes (Figure 3C). DAB staining showed that those mutants defective in SA-Signaling or biosynthesis (as represented by *pad4*) showed no H_2_O_2_ accumulation in PM-invaded epidermal cells of either –Si or +Si-treated plants (Figure 3A).

To further determine if very low leaf Si content somehow constitutively activates a SA-dependent defense mechanism, we also measured leaf SA and JA levels of the Col-0 plants before and after infection. We found that Col-0/–Si plants indeed had slightly but significantly higher total SA levels at 0 and 3 dpi compared to Col-0/+Si plants (Figure 3D), whereas there was no significant difference in JA levels between Col-0/–Si and Col-0/+Si plants (Figure 3E). Thus, together our data suggest that very low leaf Si content ectopically activates SA-dependent basal defense against powdery mildew.

### High Si-conditioned resistance is SA-independent but PAD4-dependent

To investigate if high Si-conditioned stronger resistance has the same or a distinct mechanistic basis, we first wondered if the enhanced resistance in transgenic plants of *HvLsi1*-Col-0/+Si and *CmeLsi1*-Col-0/+Si is also associated with H_2_O_2_ production. DAB staining detected H_2_O_2_ production in sporadic mesophyll cells unrelated to fungal infection and occasionally in PM-infected epidermal cells of these plants (Figures 4B, D). However, little H_2_O_2_ production was detectable in plants of the *HvLsi1*-Col-0/–Si and *CmeLsi1*-Col-0/–Si lines and these plants supported larger and more advanced mycelial networks at 5 dpi compared to their +Si-treated counterparts (Figures 4A, C). These observations suggest that very high Si content may trigger H_2_O_2_ production in mesophyll cells and potentiate H_2_O_2_ production in PM-invaded epidermal cells.

**Figure 4.**
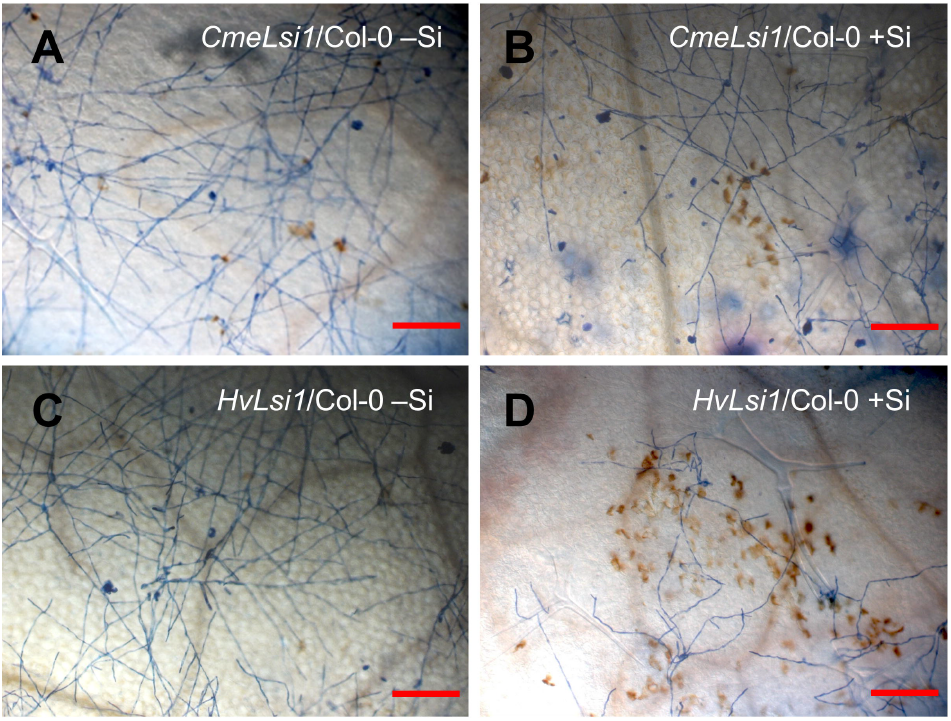
H_2_O_2_ production in sporadic mesophyll cells unrelated to fungal infection and occasionally in powdery mildew-infected epidermal cells of transgenic Col-0 treatments with Silicon. (A-D) Representative microscopic images of typical *Gc* UCSC1 fungal microcolonies grown on leaves of the indicated genotypes and treatments at 5 dpi. H_2_O_2_ reacts with 3,3-diaminobenzidine to form a reddish-brown stain, while fungal structures are stained blue by trypan blue. Scale bars = 200 μm.

Next, we introduced the same DNA constructs for expression of the two heterologous Si transporters into seven single Arabidopsis mutants (i.e. *eds1*, *pad4, sid2, coi1, ein2*), one double (*eds1pad4*) and one triple (*eds1pad4sid2*) mutant. Measurement of Si content showed that expression of *HvLsi1* or *CmeLSi1* resulted in elevation of leaf Si content in the backgrounds of all of these mutants (Figure S2), similar to that in Col-0 (Figure 2A), indicating that none of these immunity-related mutations interferes with Si uptake. We then grew plants of the above described representative transgenic lines under –Si or +Si conditions for seven weeks and then inoculated them with *Gc* UCSC1 and assessed their disease reaction phenotypes. As shown in Figure 5A&B, all transgenic lines without Si supplement exhibited disease susceptibility phenotypes as expected based on their genotypes (i.e. those SA-pathway defective mutants were more susceptible than Col-0 and those defective in JA or ET pathways). Strikingly, for the transgenic plants supplemented with Si, only plants of those genotypes that contain the *pad4* mutation, i.e. *pad4*, *eds1pad4* and *eds1pad4sid2* did not show enhanced disease resistance relative to their counterparts without Si supplement (Figure 5A-D). These results indicate that high Si content-mediated resistance is EDS1-, SA- and JA/ET-independent but PAD4-dependent.

**Figure 5.**
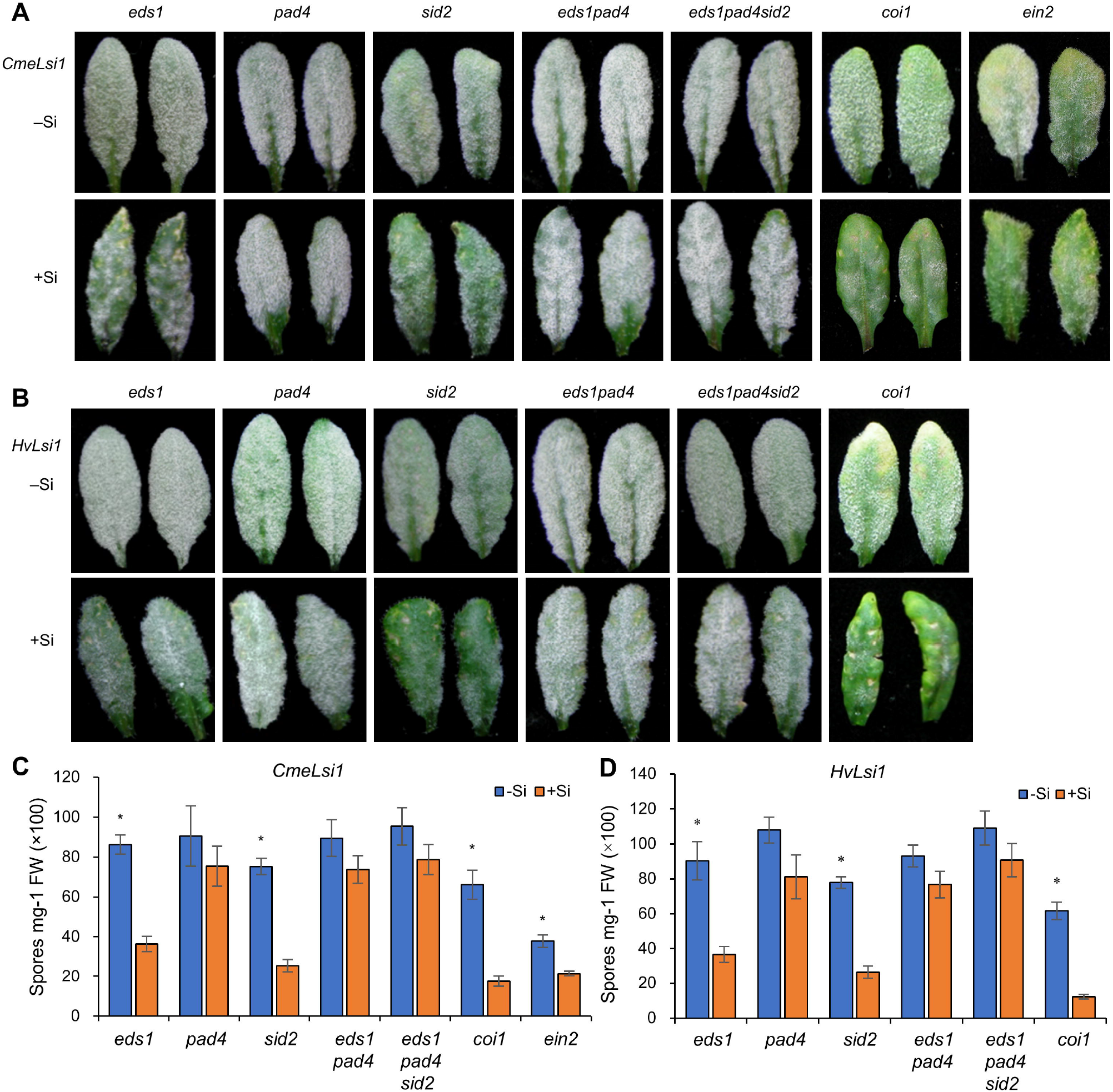
PAD4 is required for high Silicon (Si)-induced resistance against powdery mildew *Golovinomyces cichoracearum* (*Gc*) UCSC1. (A, B) Representative images of Arabidopsis leaves of the indicated mutant genotypes overexpressing *CmeLsi1* or *HvLsi1* treated with either 0 mM (–Si) or 1.0 mM (+Si) Si infected with *Gc* UCSC1 at 10 dpi. (C, D) Quantification of spore production in the indicated genotypes at 10 dpi normalized to leaf fresh weight. Bars represent standard errors and within treatments, and an asterisk on a column denotes a significant difference (*t*-test, *P<0.05).

To evaluate the molecular attributes of the PAD4-dependent mechanism activated by high Si content, we first used qRT-PCR to measure the expression levels of *PR1* in plants of *HvLsi1*-Col-0 plants with or without Si treatment at 0, 3, 5 dpi with *Gc* UCSC1. We found that *PR1* in *HvLsi1*-Col-0/+Si plants was highly expressed before powdery mildew infection (0 dpi) and had no or only slight increase at 3 or 5 dpi (Figure S3A). This observation is in agreement with H_2_O_2_ production independent of powdery mildew infection in *HvLsi1*-Col-0/+Si plants (Figure 4D). Next, we examined if high leaf Si content can increase *PAD4* expression. Interestingly, we found that although *PAD4* was induced to higher levels by powdery mildew infection at 3 and 5 dpi, no significant difference was detected between –Si and +Si plants (Figure S3B). This result suggests that high-level Si may not impact transcription of *PAD4* but rather augment certain functionality of *PAD4* via an unknown post-transcriptional mechanism, thereby activating this defense pathway. Not surprisingly, we detected no significant difference in *PDF1.2* expression between –Si and +Si plants before and after powdery mildew infection (Figure S3C), which was consistent with the observation that no significant phenotypic difference was found between transgenic lines of Col-0 and those of *coi1* (Figure 5). We also measured *PMR4* expression under different Si conditions given that the callose formation and deposition may coordinate with biological silicification in Arabidopsis (Brugiére and Exley 2017). The result showed that Si content had little impact on *PMR4* expression (Figure S3D). To further test if the PAD4-dependent, high Si-conditioned resistance is influenced by SA or JA biosynthesis, we also measured the total SA and JA levels in the transgenic –Si and +Si plants and found that levels of total SA were significantly higher in *HvLsi1*-Col-0/+Si plants compared to the plants of the same line with Si-treatment before and after powdery mildew infection (Figure S4A). As expected, the levels of total SA were significantly lower in transgenic *eds1*, *pad4*, and particularly *sid2* lines in comparison with those of transgenic Col-0 under either –Si or +Si conditions (Figure S4 A,C,E,G). Interestingly, compared with the respective –Si plants, JA levels were significantly higher in *HvLsi1*-Col-0/+Si plants before and after powdery mildew infection (Figure S4B), so were the total SA and JA levels in *HvLsi1-eds1*/+Si, *HvLsi1-pad4*/+Si, and *HvLsi1–Sid2*/+Si plants in general (except in the case of SA in the *sid2* background) (Figure S4 C, D, E, F). However, the lack of correlation between either SA or JA levels and high Si-mediates resistance suggests that increased SA or JA levels cannot explain PAD4-dependent, high Si-mediated resistance in Arabidopsis.

### Elevated Si can largely restore resistance to a non-adapted powdery mildew in an immunocompromised triple mutant

*Golovinomyces cichoracearum* (*Gc*) UMSG1 infects sow thistle (*Sonchus oleraceus*) but fails to reproduce on 25 tested Arabidopsis accessions including Col-0, even though it has largely overcome penetration resistance of Arabidopsis (Wen et al. 2010). Thus, by definition, *Gc* UMSG1 is still a non-adapted powdery mildew pathogen of Arabidopsis. However, the triple mutant *eds1pad4sid2* (in the background of Col-0) is nearly fully susceptible to *Gc* UMSG1, indicating breakdown of non-host resistance of Col-0 (Zhang et al. 2018). To test if increased Si content can compensate the loss of the non-host resistance, we inoculated plants of *HvLsi1-eds1pad4sid2* and *CmeLsi1-eds1pad4sid2* with or without Si supplement with *Gc* UMSG1. Visual examination of the infection phenotypes at 10 dpi showed that while there was no obvious difference between –Si and +Si plants of *eds1pad4sid2*, there was remarkable increased resistance (i.e. fungal mass hardly visible to the naked eye) in +Si plants but not seen in –Si plants of the same *HvLsi1- or CmeLsi1-*transgenic lines (Figure 6A). Spore counting showed that there was a ~17-fold reduction in sporulation in infected plants of the same transgenic lines treated with Si versus those without Si (Figure 6B). These results indicate that high leaf Si content can largely compensate the loss of EDS1, PAD4 and SID2 and reboot resistance to a non-adapted powdery mildew pathogen via an EDS1-, PAD4- and SID2-independent mechanism.

**Figure 6.**
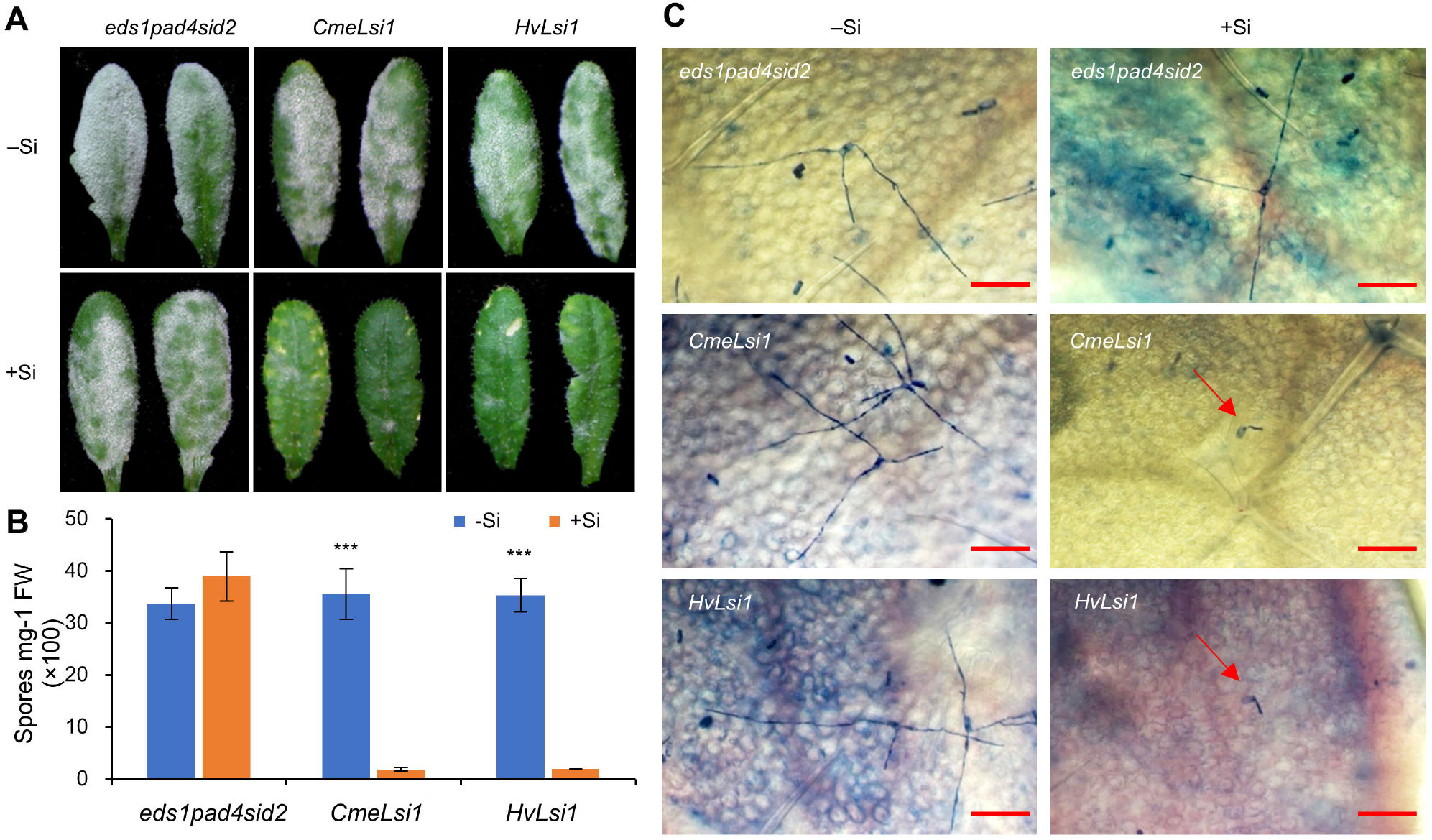
High Silicon (Si) reboots resistance to the non-adapted mildew isolate *Golovinomyces cichoracearum* (*Gc*) UMSG1 in the absence of EDS1, PAD4 and SID2. (A) Representative images of leaves of the indicated genotypes treated with either 0 mM (–Si) or 1.0 mM (+Si) Si infected with *Gc* UMSG1 at 10 dpi. (B) Quantification of spore production in the indicated genotypes at 10 dpi normalized to leaf fresh weight. Bars represent standard errors and within each genotype, and asterisks on a column denote a significant difference (t-test, ***P<0.001). (C) Representative microscopic images of typical *Gc* UMSG1 fungal microcolonies grown on leaves of the indicated genotypes and treatments at 2 dpi. Scale bars = 100 μm.

*Gc* UMSG1 is arrested after its penetration of the cell wall in leaves of Col-0 (Wen et al. 2010). To find out at what stage *Gc* UMSG1 was arrested in plants of *HvLsi1- or CmeLsi1-eds1pad4sid2*/+Si lines, we examined fungal growth in the leaves of the same transgenic lines under –Si or +Si conditions at 2 dpi and 5 dpi using Trypan Blue staining. We found that appressorial penetration of sporelings was almost completely restricted in +Si transgenic plants at 2 dpi (Figure 6C), with only rare exceptions (~1/20 sporelings) where the sporelings further differentiated limited hyphae at 5 dpi (Figure S5), some of which eventually continued to grow into sporadic colonies with sporulation at 10 dpi (Figure 6A). These observations suggest that high Si content can largely restore the penetration resistance lost in the immunocompromised Arabidopsis mutant plants against the non-adapted *Gc* UMSG1 isolate. This resistance mechanism is obviously different from that of the PAD4-dependent resistance mediated by high Si against the well-adapted *Gc* UCSC1 isolate where no gross difference in hyphal growth of sporelings at 2 dpi was observed between –Si and +Si plants (Figure S6).

## DISCUSSION

Si has long been known to increase disease resistance in plants. The suppressive effect of Si on powdery mildew was first reported in 1983 (Miyake and Takahashi 1983). However, the molecular mechanisms underlying Si-mediated resistance remain largely elusive and even controversial still today. In this study, by using the Arabidopsis-powdery mildew pathosystem, we collected genetic evidence to demonstrate that Si affects different layers of plant defense in a dosage-dependent manner, providing novel mechanistic insights into Si’s prophylactic role in plants against fungal pathogens.

A few previous studies on how Si might affect basal resistance to adapted powdery mildew using Arabidopsis Col-0 wild-types did not generate consistent results (Ghanmi et al. 2004; Fauteux et al. 2006; Vivancos et al. 2015). One contributing factor might be that different types or even batches of commercial soil for growing Arabidopsis may vary in Si content. This prompted us to use perlite as soil medium (Xiao et al. 2003) in this study to enable tighter control of Si supplement, which led to our observation that Col-0 plants with a very low leaf Si content (<2.0 mg g^−1^) displayed slightly but visually discernable and statistically significant enhanced resistance against the adapted powdery mildew *Gc* UCSC1 (Figure 1). The resistance was found to be associated with H_2_O_2_ accumulation in the invaded cells and require intact SA-signaling (Figure 3). Interestingly, a previous study also showed that oat plants deprived for Si exhibited higher phenylalanine ammonia lyase activity compared to those with normal Si supply, which led to a speculation that lack of Si may activate a compensatory mechanism resulting in better resistance to powdery mildew penetration resistance (Carver et al. 1998). Hence, we speculate that Si content below a certain threshold level (<2.0 mg g^−1^ for Arabidopsis leaves) may produce a warning signal in plant cells to activate and/or potentiate SA signaling. Considering the positive role of high Si in plant resistance against powdery mildew pathogens based on data from this and other studies (Vivancos et al. 2015, Bélanger et al. 2003), the Si dosage-impact characteristics we observed is reminiscent of the case concerning PMR4-dependent callose: Whereas deposition of callose can strengthen the papilla as a physical barrier and more extensive and faster deposition of callose by overexpression of the responsible callose synthase PMR4 confers complete resistance to powdery mildew (Ellinger et al. 2013), loss of PMR4 also results in enhanced resistance to powdery mildew (Nishimura et al. 2003). Such functional similarity between Si and callose may imply mechanistic connection between them. We thus checked if callose deposition is affected in PM-infected Col-0/–Si plants. Interestingly, we found that very low Si appeared to increase callose deposition to the papilla (Figure S7), possibly as an indirect consequence of enhanced SA-dependent defenses in Col-0/–Si plants.

Vivancos and colleagues used Arabidopsis Col-0 transgenic for wheat Si influx transporter (*TaLsi1*) to demonstrate that high Si absorbed by Arabidopsis can increase resistance to an adapted powdery mildew isolate (Vivancos et al. 2015). However, they showed that the high Si-enhanced resistance was not affected by the loss of PAD4 or SID2, which differs from our result that high Si-mediated resistance is PAD4-dependent (Figure 3). Our conclusion was inferred from the infection phenotypes of wild-type, single, double and triple *pad4*-containing transgenic lines expressing one of the two selected heterologous Si transporters using very even conidia inoculation (see Methods for details). It is possible that the discrepancy concerning PAD4 might have been caused by the differences in powdery mildew inoculation and/or growth conditions. Nevertheless, our results corroborated with their conclusion that high Si enhances SA-independent resistance to powdery mildew. The PAD4-dependence of high Si-mediated resistance is particularly interesting, because it suggests that high Si may activate or potentiate an intracellular defense mechanism beyond apoplastic obstruction (Coskun et al. 2019). Moreover, it also suggests that this PAD4-dependent mechanism is distinct from that activated by PAD4 and its partner EDS1 in SA-dependent resistance against biotrophic and hemi-biotrophic pathogens (Falk et al. 1999; Jirage et al. 1999; Feys et al. 2001). Thus, our results resemble the earlier finding that PAD4, but not EDS1, contributes to resistance of Arabidopsis to green peach aphids (GPA) (Pegadaraju et al. 2007). PAD4 has also been shown to be required for the enhanced resistance to GPA in two Arabidopsis mutants exhibiting heightened resistance against biotrophic pathogens (Louis et al. 2010; Louis et al. 2012; Lei et al. 2014). Given that Si has also been shown to increase resistance to piercing and sucking (phloem-feeding) insects including aphids (Reynolds et al. 2009), one may speculate that PAD4 also plays a similar role in Si-heightened resistance against insects. How PAD4 contributes to resistance (exerted by high Si) against pathogens and GPA (and potentially other insects when plants are supplemented with Si) via a SA-independent pathway is currently unknown. There are reports that implicate JA-dependent defense in Si-mediated resistance to pathogens (Ghareeb et al. 2011) and insects (Ye et al. 2013). In this study, we did find higher JA levels in +Si plants; however, we cannot establish a role of JA-signaling in high Si-mediated resistance because JA-signaling deficient *coi1* mutant plants transgenic for either of the two heterologous Si transporter genes under +Si conditions also showed resistance (Figure 5). Similar results were reported for Si-induced brown spot resistance (Van Bockhaven et al. 2015). The increased JA levels in all *HvLsi1*-transgenic Col-0/+Si plants may be associated with high Si-caused toxicity manifested as whitish spots in Arabidopsis leaves (Figure S8), which apparently did not affect powdery mildew infection as *pad4*-containing transgenic plants supplemented with high Si were very susceptible despite having similar whitish spots (Figure 5). Such toxicity was also observed when *TaLsi1* and *OsLsi1* were expressed in Arabidopsis using the *35S* promoter but was largely absent when a root specific promoter was used (Montpetit et al. 2012).

Apart for the unexpected finding with regard to PAD4-dependent Si-mediated resistance, another surprising observation from this study is the high Si-mediated restoration of penetration resistance in the *eds1pad4sid2* triple mutant background to *Gc* UMSG1 (Figure 6). This finding has several implications: First, Si may play an important role in blocking the entry of numerous non-adapted and probably poorly-adapted cell-wall penetrating fungal pathogens and perhaps insects. Such a role cannot be easily revealed unless an artificial pathosystem such as *eds1pad4sid2-Gc* UMSG1 is used, thus demonstrating the power of such a system. Second, that Si-mediated penetration resistance against non-adapted powdery mildew pathogens can occur in the absence of EDS1, PAD4 and SID2 implies that Si deposition to the cell wall and papillae is probably a mechanical process independent of the immunity status of the plant. Third, adapted PM, by definition, can largely overcome this physical barrier.

In summary, results from this study implicated Si in three unexpected and distinct defense mechanisms that lead to enhanced resistance to PM: (1) very low Si-triggered SA-dependent defense; (2) high Si-mediated PAD4-dependent defense; and (3) high Si-boosted penetration resistance. How to make sense of these seemingly distinct mechanisms? Information from previous reports (see below) and this study as a whole appears to support the “Si-callose synergy theory” (Brugiére and Exley 2017) which may offer a plausible explanation for our observations. First, Si is deposited to callose-rich papillae thereby playing a positive role in restricting cell-wall penetrating fungal pathogens such as powdery mildew (Carver et al. 1998; Bélanger et al. 2003; Shetty et al. 2012). Second, cell wall silicification appears to require callose and may be initiated by callose (Brugiére and Exley 2017; Kulich et al. 2018). Third, either very low Si (Figure 1) or lack of callose (due to loss of PMR4/GSL5) (Nishimura et al. 2003) triggers SA-dependent defense. Fourth, either high Si (Figure 3) or overexpression of PMR4/GSL5 (Ellinger et al. 2013) confers resistance to powdery mildew. Lastly, PAD4-dependent resistance to aphids is also associated with increased callose deposition (Rashid et al. 2017), similar to Si-mediated resistance to insects (Yang et al. 2018) or even nematodes (Zhan et al. 2018), despite that in the latter cases whether resistance is PAD4-dependent remains to be tested. Therefore, it is possible that PAD4 (but not EDS1) may play a critical role in the deposition of a basal level of PMR4-dependent callose to papillae, thus explaining the PAD4-dependence of high Si-mediated resistance against powdery mildew pathogens. Combining all the above information, we developed a schematic diagram to summarize our genetic data on Si and hypothesize that deposition of Si, along with callose, to the papilla in plant cells enhances defense against powdery mildew infection in three different scenarios (Figure S9). Future research is needed to investigate whether and how Si and callose may synergize with each other to fortify cell wall-based defense against fungal invasion, how exactly PAD4 regulates this defense mechanism, whether Si-mediated resistance to insects also requires a PAD4-regulatory node, and whether this mechanistic model is also applicable to medium- and high-Si-accumulating plants.

## MATERIALS AND METHODS

### Plant lines and growth conditions

All mutants used in this study were in the *Arabidopsis thaliana* accession Col-0 background. Mutants *eds1* (*eds1-2*), *pad4* (*pad4-1*), *sid2* (*sid2-2*), *eds1pad4* (*eds1-2pad4-1*), *eds1pad4sid2* (*eds1-2pad4-1sid2-2*), *coi1* (*coi1-1*) have been described previously by Zhang et al (2018). Mutant *ein2* (*ein2-1*) has been described by Guzman and Ecker (1990). Seeds were sown in Propagation Mix (Sun Gro Horticulture, Massachusetts) and cold treated (4 °C for 2 d), and seedlings were grown under 22 °C, 65% relative humidity, short day (10 h light at 125 μmol m^−2^ s^−1^, 14 h dark). For Si experiments, 12-days-old seedlings grown in regular soil (Propagation Mix) were transplanted into pots containing horticultural perlite (Whittemore). Plants were irrigated with nutrient solution prepared with deionized pure water. The nutrient solution was prepared based on Boursiac et al. (2010) with some modifications (pH 5.8; 1.5 mM Ca(NO_3_)_2_·4H_2_O, 1.25 mM KNO_3_, 0.5 mM KH_2_PO_4_, 0.75 mM MgSO_4_·7H_2_O, 0.046 mM H_3_BO_3_, 9.55 μM MnSO_4_·4H_2_O, 0.77 μM ZnSO_4_·7H_2_O, 0.32 μM CuSO_4_·5H_2_O, 0.016 μM (NH_4_)_6_Mo_7_O_24_·4H_2_O, 0.054 mM EDTA-bisodium salt). When plants were 6 weeks old, half of them were irrigated with nutrient solution containing 1.0 mM Na_2_SiO_3_·9H_2_O (+Si), the other half were irrigated with nutrient solution containing 1.0 mM Na_2_SO_4_ (–Si).

### DNA constructs and generation of transgenic lines

To make Arabidopsis absorb more Si from soil medium, we generated stable transgenic lines of Col-0 wild-type and various mutants that express *HvLsi1*(a barley Si transporter gene; GenBank accession LOC100301576) or *CmeLsi1*(a muskmelon Si transporter gene; GenBank accession LOC103487002) from the *35S* promoter. Briefly, total RNA was extracted from roots of barley (*Hordeum vulgare* L.) roots or muskmelon (*Cucumis melo* L.) using the TRIZOL reagent (Invitrogen) according to the manufacturer’s recommendations and stored at −80°C until use. First strand cDNAs were prepared from 1 μg of total RNA using a reverse transcriptase (Superscript III, Invitrogen) and Oligo(dT)18 primers. The coding sequences of *HvLsi1* and *CmeLsi1* were amplified with the Q5 DNA polymerase (New England Biolabs, M0491L) with appropriate primer pairs (see Supplemental Table S1). The DNA fragments were cloned into pENTR/D-TOPO (Thermo Fisher Scientific Inc.) and shuttled to the Gateway Compatible binary vector pEarleyGate100. After DNA sequence confirmation, the constructs were introduced into Arabidopsis plants via *Agrobacterium*-mediated transformation using the *A. tumefaciens* strain GV3101. At least 20 independent T1 transgenics were obtained for each DNA construct/genotype combination. T2 progenies (24 plants) of at least ten T1 lines were grown under –Si and +Si conditions and inoculated with powdery mildew to visually assess the infection phenotypes. T3 generations derived from three T1 independent lines were used to confirm the infection phenotypes, and one representative homozygous T3 line for each genotype was used for comparative and quantitative analysis with other relevant genotypes.

### Pathogen infection and quantification of disease phenotypes

Adapted powdery mildew isolate *Golovinomyces cichoracearum* (*Gc*) UCSC1 was maintained on *pad4* plants and the non-adapted isolate *Gc* UMSG1 was maintained on sow thistle plants (Wen et al. 2010). Inoculation and visual scoring of disease reaction phenotypes and spore quantification were done as previously described (Zhang et al. 2018). Briefly, for quantification of disease susceptibility, five or six duplicate leaf samples (each consisting of ~120 mg leaves) collected from 12 plants of each representative T3 line at 10 dpi were used to quantify the level of sporulation. A spore suspension (or 10x dilution if the genotype was very susceptible) of each sample, which was made by vortexing the leaves in a 50 ml falcon tube containing 10 ml of H_2_O + 0.02% Silwet L-77 (Lehle seeds, USA) for one minute, was used for spore counting using Luna™ Automated Cell Counter (Logos biosysems). Spore counts were normalized to the fresh weight of the corresponding leaf samples. All infection trials with T3 generations were repeated three times with similar results, and data from one experiment were presented.

### Detection of H_2_O_2_, callose deposition and fungal structures

The detection of H_2_O_2_ accumulation in leaf tissues by 3,3-diaminobenzidine (DAB) staining was modified from Thordal-Christensen et al. (1997). Inoculated leaves were excised at the base of the petiole, placed in 1 mg ml^−1^ DAB (Sigma), and incubated for 6 h at 25°C with illumination. Fungal structures in inoculated leaves were visualized with 0.25% Trypan blue staining solution (Xiao et al. 2003). Callose deposition at the fungal penetration sites (i.e. papillae) and around the haustorium was detected by aniline blue staining. Leaves were cleared in a solution containing ethanol, water, acetic acid, and glycerol (8:1:1:1) for 48 hours at 37 °C with one change of the solution. The cleared leaves are then stained with 0.01% aniline blue in an aqueous solution containing 150 mM KH_2_PO_4_ (PH 9.5) for 4 hours. Callose deposition was visualized by fluorescence microscopy.

### Measurement of Si content in Arabidopsis leaves

Leaf Si content was determined ten days after treatment with +Si or –Si nutrient solution using seven or eight-week-old plants. All rosette leaves from five +Si or –Si plants per sample were oven dried at 65°C for 72 h, and ground into a fine powder using a mortar and pestle before measurement of Si concentration by colorimetric analysis using 0.1 g alkali-digested leaf tissue powder (Frantz et al. 2008). Three duplicated samples were processed for each genotype-treatment combination and Si content was calculated, adjusted for dry weight of leaf tissues used, and presented as mg Si dioxide per gram dry matter.

### qRT-PCR analysis

Gene expression was measured by qRT-PCR according to Zhang et al. 2018 with minor modification. Five duplicate leaf samples (~100 mg each) per genotype-treatment were collected at 0, 3 and 5 dpi with *Gc* UCSC1. The transcript levels of the target genes were normalized to that of UBC9 (Ubiquitin conjugating enzyme 9, AT4G27960). Data were analyzed by using the comparative ΔΔCt method (Livak and Schmittgen 2001). Primers are listed in Supplemental Table S1.

### Measurement of levels of SA and JA

Five leaf samples (~150 mg each) per genotype-treatment were harvested at 0, 3 and 5 dpi with *Gc* UCSC1 for determining levels of both SA and JA as previously described (Floková et al. 2014), with some modifications. For each experiment, detection was performed with three biological replicates per treatment. The leaf tissues in a 1.5 ml tube were added with 1 ml of 50% ethanol containing the internal standards (vanillic acid and dihydro jasmonic acid), four steel balls (diameter 5 mm), and then shaken for 5×1 minute in a TissueLyser II (QIAGEN) at 25Hz with one min pause between every min before centrifuged at 20,000g for 10 mins. The supernatant was analyzed with a Waters Acquity UPLC system equipped with a Waters LCT Premiere XE ESI-TOF mass spectrometer. The detailed method is described in Methods S1.

## ACKNOWLEDGMENTS

We thank Franker Coker for maintaining the plant growth facilities. This work was in part supported by a National Science Foundation grant (IOS-1457033) and a Maryland Horticultural Society grant (2017) to S. X., and a scholarship from the China Scholarship Council to L. W.

## AUTHOR CONTRIBUTIONS

X. D., L.W. and S. X. planned and designed the research. L. W., M. D., Y. W., Q. Z., L. H., E. E. and J. F P. performed experiments and analyzed data. L.W. and S. X. wrote the manuscript.

## SUPPORTING INFORMATION

**Methods S1.** SA and JA levels were detected with a Waters Acquity UPLC system equipped with a Waters LCT Premiere XE ESI-TOF mass spectrometer.

**Figure S1.** *Gc* UCSC1 infection phenotypes of Col-0 grown in normal soil treated with either 0 mM or 1.7 mM Si.

**Figure S2.** Silicon content in leaves of different Arabidopsis mutant and transgenic mutant lines.

**Figure S3.** Expression levels of defense-related genes in transgenic Col-0 overexpressing *HvLsi1*.

**Figure S4.** Levels of total SA and JA in Arabidopsis plants overexpressing *HvLsi1*.

**Figure S5.** Microscopic images showing fungal microcolonies grown on leaves at 5 dpi.

**Figure S6.** Microscopic images showing fungal microcolonies grown on leaves at 2 dpi.

**Figure S7.** Callose deposition in Arabidopsis Col-0 leaves inoculated with *G. cichoracearum* UCSC1.

**Figure S8.** Leaf phenotypes of Col-0 and *pad4-1* and their transgenic plants overexpressing the indicated Si transporter grown in perlite without Si or with 1.0 mM Si.

**Figure S9.** Schematic diagram describing the three scenarios where very low Silicon (Si) or high Si confers enhanced resistance to powdery mildew, and the potential synergistic action between Si and callose.

**Table S1.** Primers used in this work.

## REFERENCES

Bélanger RR, Benhamou N, Menzies JG (2003) Cytological evidence of an active role of silicon in wheat resistance to powdery mildew (*Blumeria graminis* f. sp. *tritici*). Phytopathology 93: 402–412.

Boursiac Y, Lee SM, Romanowsky S, Blank R, Sladek C, Chung WS, Harper JF (2010) Disruption of the vacuolar calcium-ATPases in Arabidopsis results in the activation of a salicylic acid-dependent programmed cell death pathway. Plant Physiol 154: 1158–1171.

Brugiére T, Exley C (2017) Callose-associated silica deposition in Arabidopsis. J Trace Elem Med Biol 39: 86–90.

Brunings AM, Datnoff LE, Ma JF, Mitani N, Nagamura Y, Rathinasabapathi B, Kirst M (2009) Differential gene expression of rice in response to silicon and rice blast fungus *Magnaporthe oryzae*. Ann Appl Biol 155: 161–170.

Carver TLW, Robbins MP, Thomas BJ, Troth K, Raistrick N, Zeyen R J (1998) Silicon deprivation enhances localized autofluorescent responses and phenylalanine ammonia-lyase activity in oat attacked by *Blumeria graminis*. PMPP 52: 245–257.

Chain F, Côté-Beaulieu C, Belzile F, Menzies JG, Bélanger RR (2009) A comprehensive transcriptomic analysis of the effect of silicon on wheat plants under control and pathogen stress conditions. MPMI 22: 1323–1330.

Chiba Y, Mitani N, Yamaji N, Ma JF (2009) HvLsi1 is a silicon influx transporter in barley. Plant J 57: 810–818.

Coskun D, Deshmukh R, Sonah H, Menzies JG, Reynolds O, Ma JF, Kronzucker HJ, Bélanger RR (2019) The controversies of silicon’s role in plant biology. New Phytol 221: 67–85.

Ellinger D, Naumann M, Falter C, Zwikowics C, Jamrow T, Manisseri C, Somerville SC, Voigt CA (2013) Elevated early callose deposition results in complete penetration resistance to powdery mildew in Arabidopsis. Plant Physiol 16: 1433–1444.

Epstein E (1994) The anomaly of silicon in plant biology. Proc Natl Acad Sci USA 91: 11–17.

Falk A, Feys BJ, Frost LN, Jones JDG, Daniels MJ, Parker JE (1999) EDS1, an essential component of R gene-mediated disease resistance in Arabidopsis has homology to eukaryotic lipases. Proc Natl Acad Sci USA 96: 3292–3297.

Fauteux F, Chain F, Belzile F, Menzies JG, Bélanger RR (2006) The protective role of silicon in the Arabidopsis-powdery mildew pathosystem. Proc Natl Acad Sci USA 103: 17554–17559.

Feys BJ, Moisan LJ, Newman MA, Parker JE (2001) Direct interaction between the Arabidopsis disease resistance signaling proteins, EDS1 and PAD4. EMBO J 20: 5400–5411.

Floková K, Tarkowská D, Miersch O, Strnad M, Wasternack C, Novák O (2014) UHPLC – MS/MS based target profiling of stress-induced phytohormones. Phytochemistry 105: 147–157.

Frantz JM, Locke JC, Datnoff L, Omer M, Widrig A, Sturtz D, Horst L, Krause CR (2008) Detection, distribution, and quantification of silicon in floricultural crops utilizing three distinct analytical methods. Commun Soil Sci Plant Anal 39: 2734–2751.

Ghanmi D, McNally DJ, Benhamou N, Menzies JG, & Bélanger RR (2004) Powdery mildew of Arabidopsis thaliana: a pathosystem for exploring the role of silicon in plant-microbe interactions. PMPP 64: 189–199.

Ghareeb H, Bozsó Z, Ott PG, Repenning C, Stahl F, Wydra K (2011) Transcriptome of silicon-induced resistance against Ralstonia solanacearum in the silicon non-accumulator tomato implicates priming effect. PMPP 75: 83–89.

Guzman P, Ecker JR (1990) Exploiting the triple response of Arabidopsis to identify ethylene-related mutants. Plant Cell 2: 513–523.

Jirage D, Tootle TL, Reuber TL, Frost LN, Feys BJ, Parker JE, Ausubel FM, Glazebrook J (1999) *Arabidopsis thaliana* PAD4 encodes a lipase-like gene that is important for salicylic acid signaling. Proc Natl Acad Sci USA 96: 13583–13588.

Kulich I, Vojtíková Z, Sabol P, Ortmannová J, Neděla V, Tihlaříková E, & Žárský V (2018) Exocyst subunit EXO70H4 has a specific role in callose synthase secretion and silica accumulation. Plant Physiol 176: 2040–2051.

Lei J, Finlayson SA, Salzman RA, Shan L, Zhu-Salzman K (2014) BOTRYTIS-INDUCED KINASE1 modulates Arabidopsis resistance to green peach aphids via PHYTOALEXIN DEFICIENT4. Plant Physiol 165:1657–1670.

Livak KJ, Schmittgen TD (2001) Analysis of relative gene expression data using real-time quantitative PCR and the 2^-ΔΔCT^ method. Methods 25: 402–408.

Louis J, Gobbato E, Mondal HA, Feys BJ, Parker JE, Shah J (2012) Discrimination of Arabidopsis PAD4 activities in defence against green peach aphid and pathogens. Plant Physiol 158:1860–1872.

Louis J, Leung Q, Pegadaraju V, Reese J, Shah, J (2010) PAD4-dependent antibiosis contributes to the ssi2-conferred hyper-resistance to the green peach aphid. MPMI 23: 618–627.

Ma JF (2010) Silicon transporters in higher plants. In: Jahn TP, Bienert GP, eds. MIPs and their role in the exchange of metalloids. Springer, New York. pp. 99–109.

Ma JF, Tamai K, Yamaji N, Mitani N, Konishi S, Katsuhara M, Ishiguro M, Murata Y, Yano M (2006) A silicon transporter in rice. Nature 440: 688.

Menzies J, Bowen P, Ehret D, Glass AD (1992) Foliar applications of potassium silicate reduce severity of powdery mildew on cucumber, muskmelon, and zucchini squash. J Am Soc Hortic Sci 117: 902–905.

Meyer D, Pajonk S, Micali C, O’Connell R, Schulze-Lefert P (2009) Extracellular transport and integration of plant secretory proteins into pathogen-induced cell wall compartments. Plant J 57: 986–999

Mitani N, Yamaji N, Ago Y, Iwasaki K, Ma, JF (2011) Isolation and functional characterization of an influx silicon transporter in two pumpkin cultivars contrasting in silicon accumulation. Plant J 66: 231–240.

Miyake Y, Takahashi E (1983) Effect of silicon on the growth of solution-cultured cucumber plant. Soil Sci Plant Nutr 29: 71–83.

Montpetit J, Vivancos J, Mitani-Ueno N, Yamaji N, Rémus-Borel W, Belzile F, Ma JF, Bélanger RR (2012) Cloning, functional characterization and heterologous expression of TaLsi1, a wheat silicon transporter gene. Plant Mol Biol 79: 35–46.

Nishimura MT, Stein M, Hou BH, Vogel JP, Edwards H, Somerville SC (2003) Loss of a callose synthase results in salicylic acid-dependent disease resistance. Science 301: 969–972.

Pegadaraju V, Louis J, Singh V, Reese JC, Bautor J, Feys BJ, Cook G, Parker JE, Shah J (2007) Phloem-based resistance to green peach aphid is controlled by Arabidopsis PHYTOALEXIN DEFICIENT4 without its signaling partner ENHANCED DISEASE SUSCEPTIBILITY1. Plant J 52: 332–341.

Pieterse CMJ, Leon-Reyes A, Van der Ent S, Van Wees SC (2009) Networking by small-molecule hormones in plant immunity. Nat Chem Biol 5: 308.

Rafi MM, Epstein E, Falk RH (1997) Silicon deprivation causes physical abnormalities in wheat (*Triticum aestivum* L). J Plant Physiol 151: 497–501.

Rashid MH, Khan A, Hossain MT, Chung YR (2017) Induction of Systemic Resistance against Aphids by Endophytic Bacillus velezensis YC7010 via Expressing PHYTOALEXIN DEFICIENT4 in Arabidopsis. Front Plant Sci 8: 211.

Reynolds OL, Keeping M, Meyer J (2009) Silicon-augmented resistance of plants to herbivorous insects: a review. Ann Appl Biol 155: 171–186.

Shetty R, Fretté X, Jensen B, Shetty NP, Jensen JD, Jørgensen HJL, Newman MA, Christensen LP (2011) Silicon-induced changes in antifungal phenolic acids, flavonoids, and key phenylpropanoid pathway genes during the interaction between miniature roses and the biotrophic pathogen *Podosphaera pannosa*. Plant Physiol 157: 2194–2205.

Shetty R, Jensen B, Shetty NP, Hansen M, Hansen CW, Starkey KR, Jørgensen HJL (2012) Silicon induced resistance against powdery mildew of roses caused by *Podosphaera pannosa*. Plant Pathol 61: 120–131.

Takahashi E, Ma JF, Miyake Y (1990) The possibility of silicon as an essential element for higher plants. Comments on Agricultural and Food Chemistry 2: 99–102.

Thordal-Christensen H, Zhang Z, Wei Y, Collinge DB (1997) Subcellular localization of H_2_O_2_ in plants. H_2_O_2_ accumulation in papillae and hypersensitive response during the barley-powdery mildew interaction. Plant J 11: 1187–1194.

Van Bockhaven J, De Vleesschauwer D, Höfte M (2012) Towards establishing broad-spectrum disease resistance in plants: silicon leads the way. J Exp Bot 64: 1281–1293.

Van Bockhaven J, Spíchal L, Novák O, Strnad M, Asano T, Kikuchi S, Höfte M, De Vleesschauwer D (2015) Silicon induces resistance to the brown spot fungus Cochliobolus miyabeanus by preventing the pathogen from hijacking the rice ethylene pathway. New Phytol 206: 761–773.

Vivancos J, Labbé C, Menzies J G, Bélanger RR (2015) Silicon-mediated resistance of Arabidopsis against powdery mildew involves mechanisms other than the salicylic acid (SA)-dependent defence pathway. Mol Plant Pathol 16: 572–582.

Wei Y, Zhang Z, Andersen CH, Schmelzer E, Gregersen PL, Collinge DB, Smedegaard-Petersen V, Thordal-Christensen H (1998) An epidermis/papilla-specific oxalate oxidase-like protein in the defence response of barley attacked by the powdery mildew fungus. Plant Mol Biol 36: 101–112.

Wen Y, Wang W, Feng J, Luo, MC, Tsuda K, Katagiri F, Bauchan G, Xiao S (2010) Identification and utilization of a sow thistle powdery mildew as a poorly adapted pathogen to dissect post-invasion non-host resistance mechanisms in Arabidopsis. J Exp Bot 62: 2117–2129.

Wiese J, Wiese H, Schwartz J, Schubert S (2005) Osmotic stress and silicon act additively in enhancing pathogen resistance in barley against barley powdery mildew. J Plant Nutr Soil Sci 168: 269–274.

Xiao S, Brown S, Patrick E, Brearley C, Turner JG (2003) Enhanced transcription of the Arabidopsis disease resistance genes RPW8. 1 and RPW8. 2 via a salicylic acid-dependent amplification circuit is required for hypersensitive cell death. Plant Cell 15: 33–45.

Yang L, Li P, Li F, Ali S, Sun X, Hou M (2018) Silicon amendment to rice plants contributes to reduced feeding in a phloem-sucking insect through modulation of callose deposition. Ecol Evol 8: 631–637.

Ye M, Song Y, Long J, Wang R, Baerson SR, Pan Z, Zhu-Salzman K, Xie J, Cai K, Luo S, et al (2013) Priming of jasmonate-mediated antiherbivore defense responses in rice by silicon. Proc Natl Acad Sci USA 110: E3631–E3639.

Yoshida S (1965) Chemical aspects of the role of silicon in physiology of the rice plant. Bull Nat Inst Agr Sci 15: 18–58.

Zhan LP, Peng DL, Wang XL, Kong LA, Peng H, Liu SM, Liu Y, Huang WK (2018) Priming effect of root-applied silicon on the enhancement of induced resistance to the root-knot nematode *Meloidogyne graminicola* in rice. BMC Plant Biol 18: 50.

Zhang Q, Berkey R, Blakeslee JJ, Lin J, Ma X, King H, Liddle A, Guo L, Munnik T, Wang X, et al(2018) Arabidopsis phospholipase Dα1 and Dδ oppositely modulate EDS1-and SA-independent basal resistance against adapted powdery mildew. J Exp Bot 69: 3675–3688.

