## Supplemental S1-S9+ MethodS1+Table S1 for "Silicon modulates multi-layered defense against powdery mildew in Arabidopsis"

### Supplementary information

#### Method S1. making nutrient solutions with Silicon (+Si) and without Silicon (–Si)

To prepare +Si nutrient solution, first, 1.0 mM $Na_2SiO_3 \cdot 9H_2O$ was made with deionized pure water, with its pH value adjusted to 7.0 to avoid calcium precipitation at later steps. Then, other nutrient elements (i.e. 1.5 mM $Ca(NO_3)_2 \cdot 4H_2O$, 1.25 mM $KNO_3$, 0.5 mM $KH_2PO_4$, 0.75 mM $MgSO_4 \cdot 7H_2O$, 0.046 mM $H_3BO_3$, 9.55 µM $MnSO_4 \cdot 4H_2O$, 0.77 µM $ZnSO_4 \cdot 7H_2O$, 0.32 µM $CuSO_4 \cdot 5H_2O$, 0.016 µM $(NH_4)_6Mo_7O_{24} \cdot 4H_2O$, 0.054 mM EDTA-bisodium salt) were added one by one sequentially according to the nutrient formulation. At last, the pH value of the solution was adjusted to 5.8 with 2M $H_2SO_4$. To prepare –Si nutrient solution, 1.0 mM $Na_2SO_4$ was made with deionized pure water. Then, the same other nutrient elements were added sequentially before adjusting the pH value to 5.8 with 2M NaOH.

#### Methods S2. SA and JA levels were detected with a Waters Acquity UPLC system equipped with a Waters LCT Premiere XE ESI-TOF mass spectrometer

Four microliter injections of leaf tissue supernatant were made and phytohormones were separated using a Waters HSS C18 column (2.1 × 100 mm; 1.7 um particle size). A binary solvent system consisting of 0.1% formic acid (solvent A) and 0.1% formic acid in acetonitrile (solvent B) was used. Gradient conditions were: 0 - 1 minute, 20% solvent B; 1 - 3.5 minutes, a linear gradient to 90% solvent B; 3.5 - 4 minutes, 90% solvent B. The flow rate was 0.35 ml $min^{-1}$. Phytohormones were quantified via mass spectrometry by integrating extracted ion chromatogram peak areas for compounds of interest. Compounds were detected in negative ion mode using the following parameters: Capilliary voltage 2150 V, cone voltage 20 V, desolvation temperature 275°C, desolvation gas flow 250 L $h^{-1}$, cone gas flow 15 L $h^{-1}$, source temperature 115°C. Standard curves were constructed using authentic phytohormone standards.

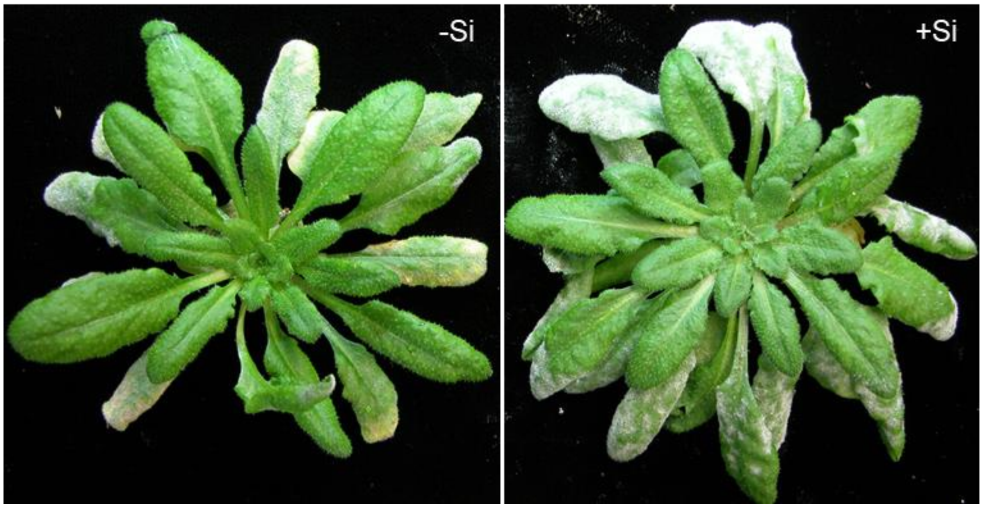


**Figure S1.** ***Gc*** **UCSC1 infection phenotypes of Col-0 grown in normal soil treated with either 0 mM or 1.7 mM Si**

Plants of Arabidopsis accession Col-0 grown in soil irrigated with 1.7 mM Silicon (Si) were slightly more susceptible than those without Si treatment to powdery mildew isolate *Golovinomyces cichoracearum* UCSC1. Two representative plants were shown. Pictures were taken at 10 dpi.

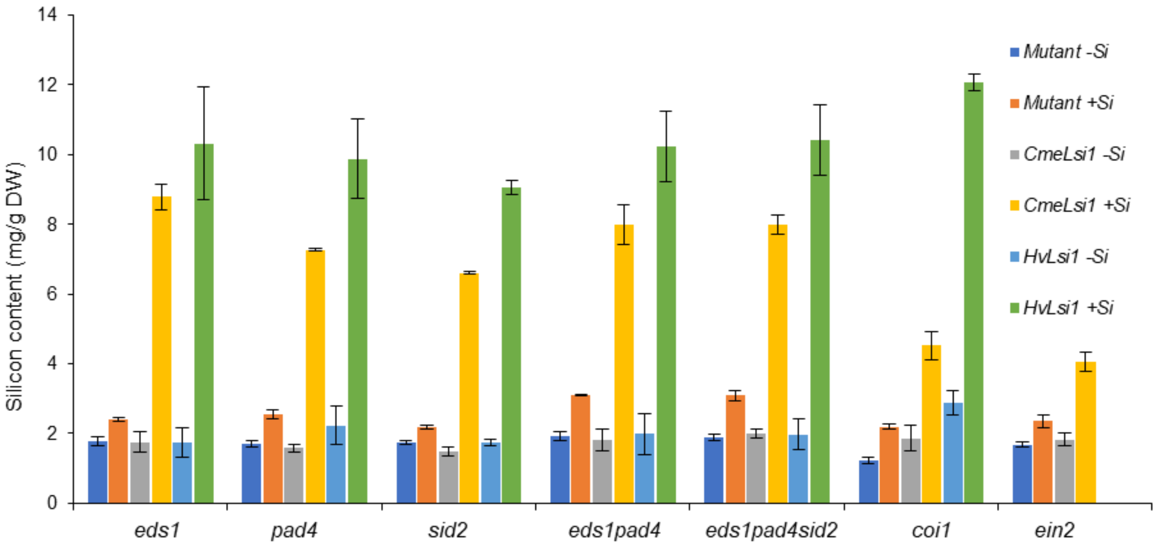


**Figure S2. Silicon content in leaves of different Arabidopsis mutant and transgenic mutant lines**

Expression of either *CmeLsi1* or *HvLsi1* from the *35S* promoter in Arabidopsis mutants defective in defense signaling resulted in elevation of leaf Silicon (Si) content. The indicated Arabidopsis mutants (indicated by "*Mutant*") and their respective transgenic lines (indicated by "*CmeLsi1*" or "*HvLsi1*" were treated with either 0 mM (–Si) or 1.0 mM (+Si) Si. The leaf Si content was normalized to leaf dry weight. Bars represent standard errors.

Click here to enter text.

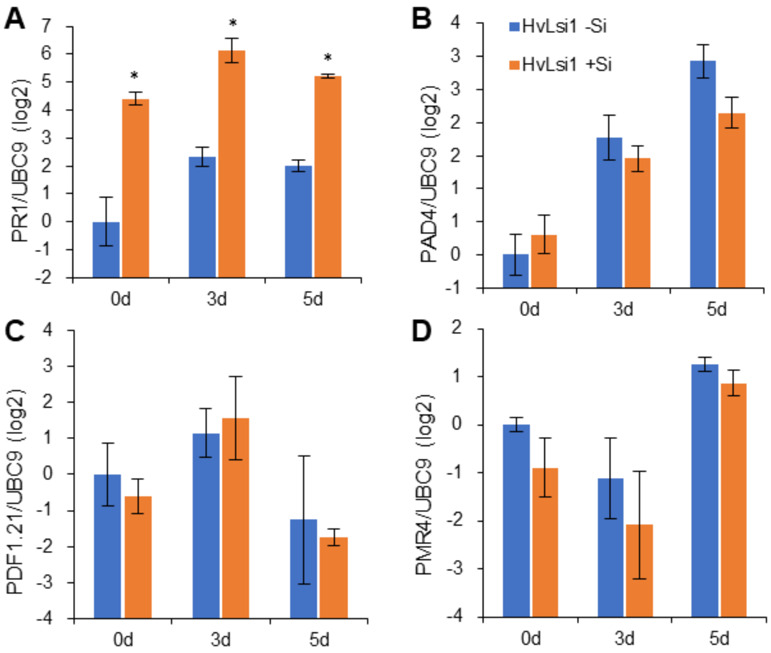


**Figure S3. Expression levels of defense-related genes in transgenic Col-0 overexpressing *HvLsi1***

Expression levels of genes encoding Pathogenesis-Related protein 1 (*PR1*, A), Phytoalexin Deficient 4 (*PAD4*, B), Plant Defensin 1.2 (*PDF1.2*, C) and Powdery Mildew Resistant 4 (*PMR4*, D) in Col-0 plants transgenic for 35S-*HvLsi1*. Plants were grown in perlite treated with either 0 mM (–Si) or 1.0 mM (+Si) Silicon and then inoculated with *Golovinomyces cichoracearum* UCSC1. Total RNA was prepared from inoculated leaves at 0, 3 and 5 dpi. Bars represent standard errors, and an asterisk denotes a significant difference between the “–Si” and “+Si” treatments (Student *t*-test, *P<0.05).

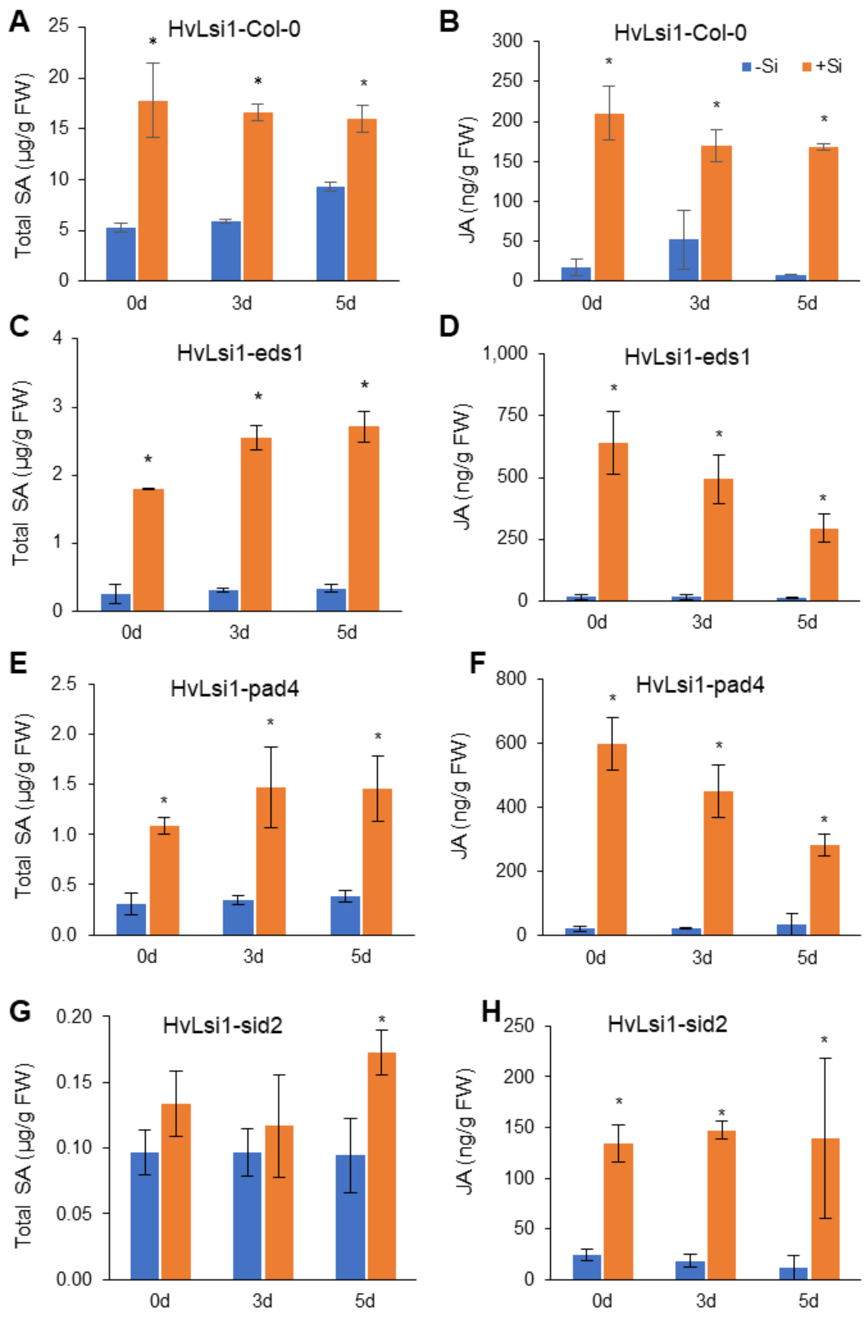


**Figure S4. Levels of total SA and JA in Arabidopsis plants overexpressing *HvLsi1***

Levels of total SA and JA in plants of the indicated genotypes transgenic for *35S-HvLsi1* treated with either 0 mM (–Si) or 1.0 mM (+Si) Si were measured at 0, 3 and 5 dpi with *Golovinomyces cichoracearum* UCSC1. Bars represent standard errors and an asterisk denotes a significant difference between the "–Si" and "+Si" treatments (Student *t*-test, *P<0.05).

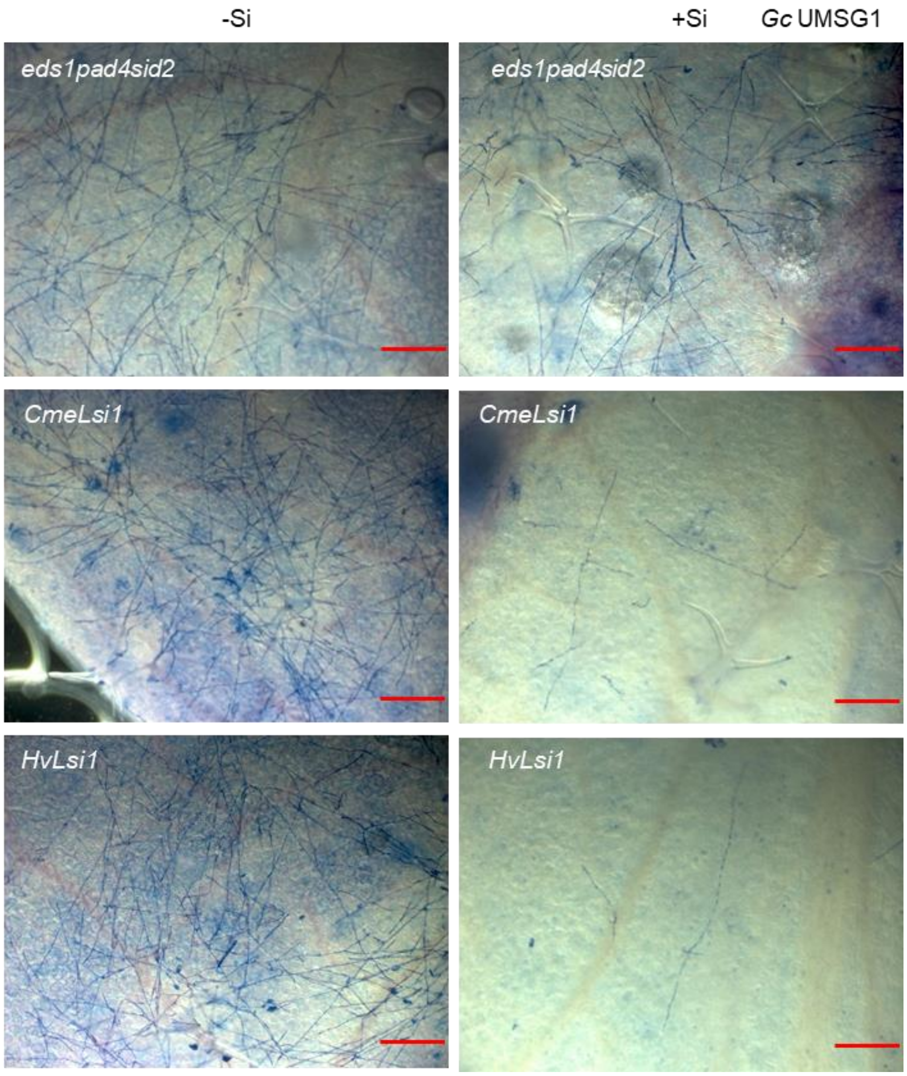


**Figure S5. Microscopic images showing fungal microcolonies grown on leaves at 5 dpi**

Plants of *eds1pad4sid2* and their transgenic lines expressing *CmeLsi1* or *HvLsi1* were treated with 0 mM (–Si) or 1.0 mM (+Si) Si and inoculated with *G. cichoracearum* (*Gc*) UMSG1. Inoculated leaves were subjected to Trypan Blue staining at 5 dpi. Representative fungal microcolonies were shown. Scale bars=200 μm.

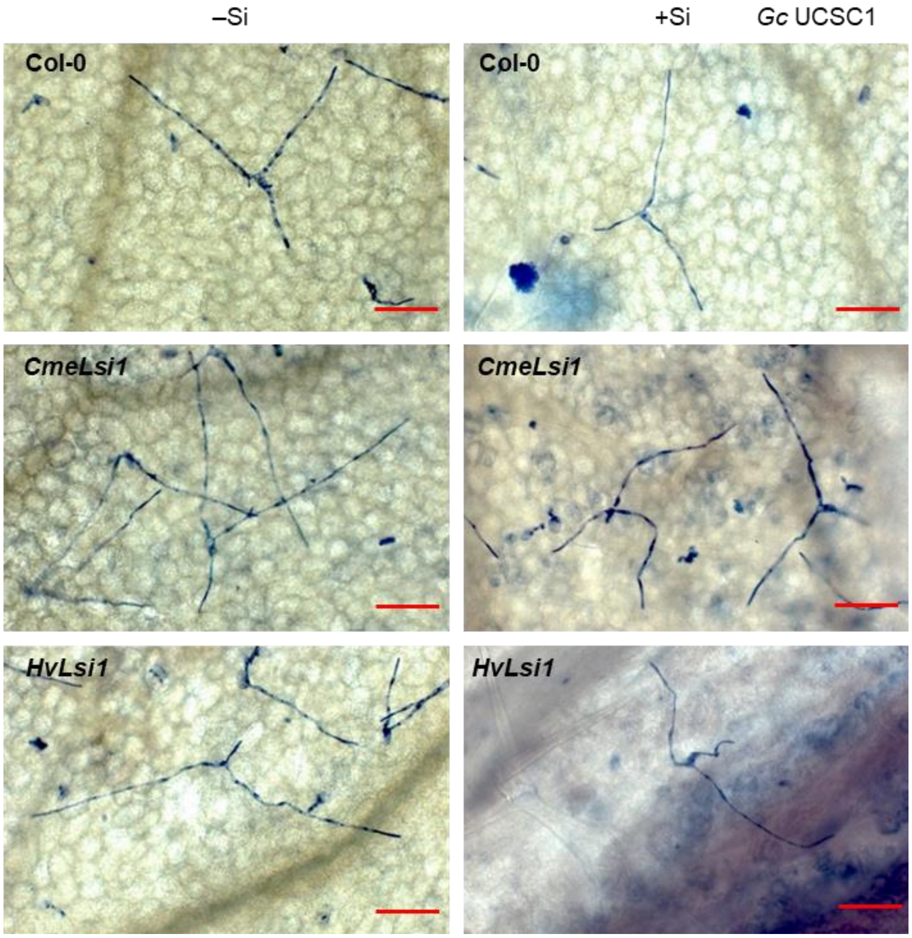


**Figure S6. Microscopic images showing fungal microcolonies grown on leaves at 2 dpi**

High Silicon (Si) did not seem to affect early infection of *G. cichoracearum* (*Gc*) UCSC1 in Col-0 plants or Col-0 transgenic for *35S-CmeLsi1* or 35S-*HvLsi1*. Plants were treated with 0 mM (–Si) or 1.0 mM (+Si) Si and inoculated with *Gc* UCSC1. Inoculated leaves were subjected to Trypan Blue staining at 2 dpi. Representative fungal microcolonies were shown. Scale bars=100 µm.

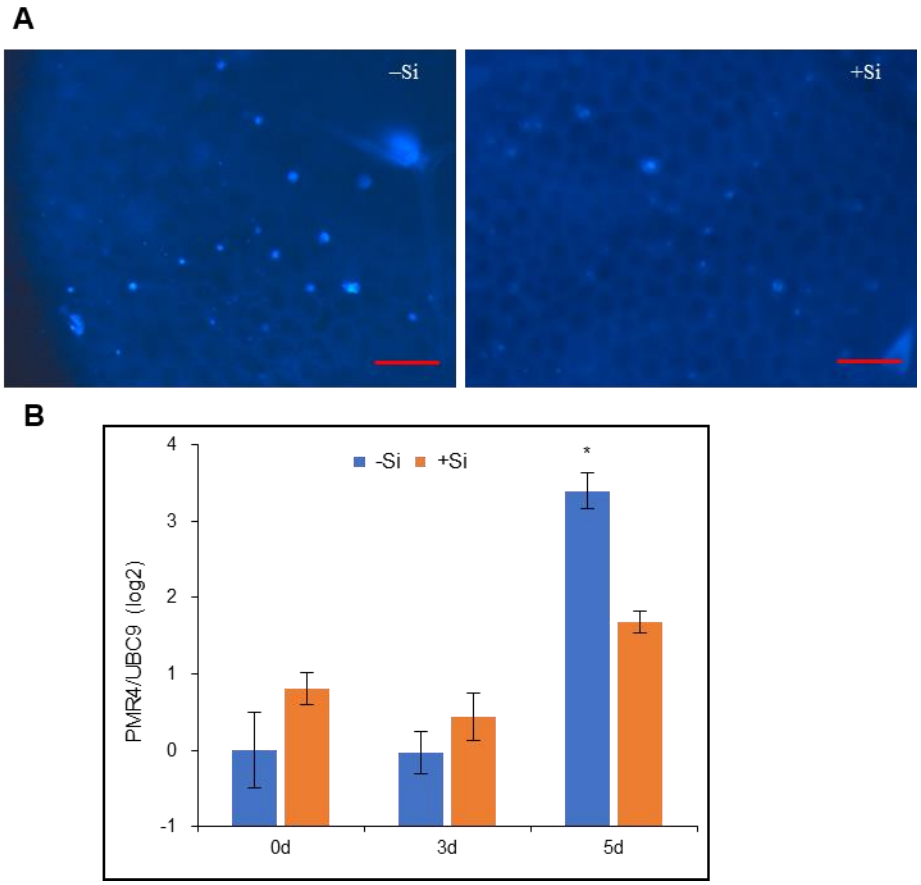


**Figure S7. Callose deposition in Arabidopsis Col-0 leaves inoculated with *G. cichoracearum* UCSC1**

(A) Callose deposition in *G. cichoracearum* UCSC1-inoculated leaves of Col-0 treated with either 0 mM (–Si) or 1.0 mM (+Si) Si at 5 dpi. Leaves were subjected to Trypan Blue staining. Callose deposition is indicated by arrowheads. Scale bars=200 µm. (B) Expression of *PMR4* in Col-0 plants used in (A) at 0, 3 and 5 dpi. Bars represent standard errors, and an asterisk denotes significant difference between the "–Si" and "+Si" treatments (Student *t*-test, *P<0.05).

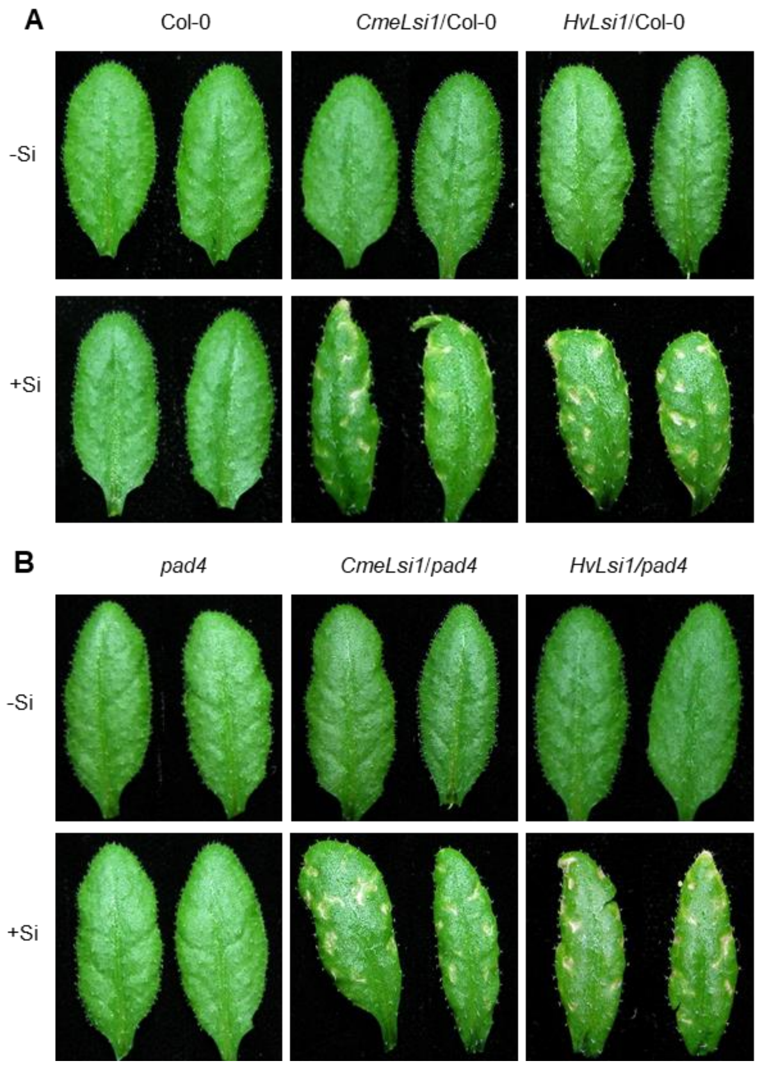


**Figure S8. Leaf phenotypes of Col-0 and *pad4-1* and their transgenic plants overexpressing the indicated Si transporter grown in perlite without Si or with 1.0 mM Si**

The toxicity was shown by whitish spots in mature leaves of the indicated plants treated with 1.0 mM Si (+Si) for two weeks. No whitish spots were observed in plants of the same transgenic lines grown with 0 mM Si (–Si) or non-transgenic Col-0 plants grown with either 0 mM or 1.0 mM Si.

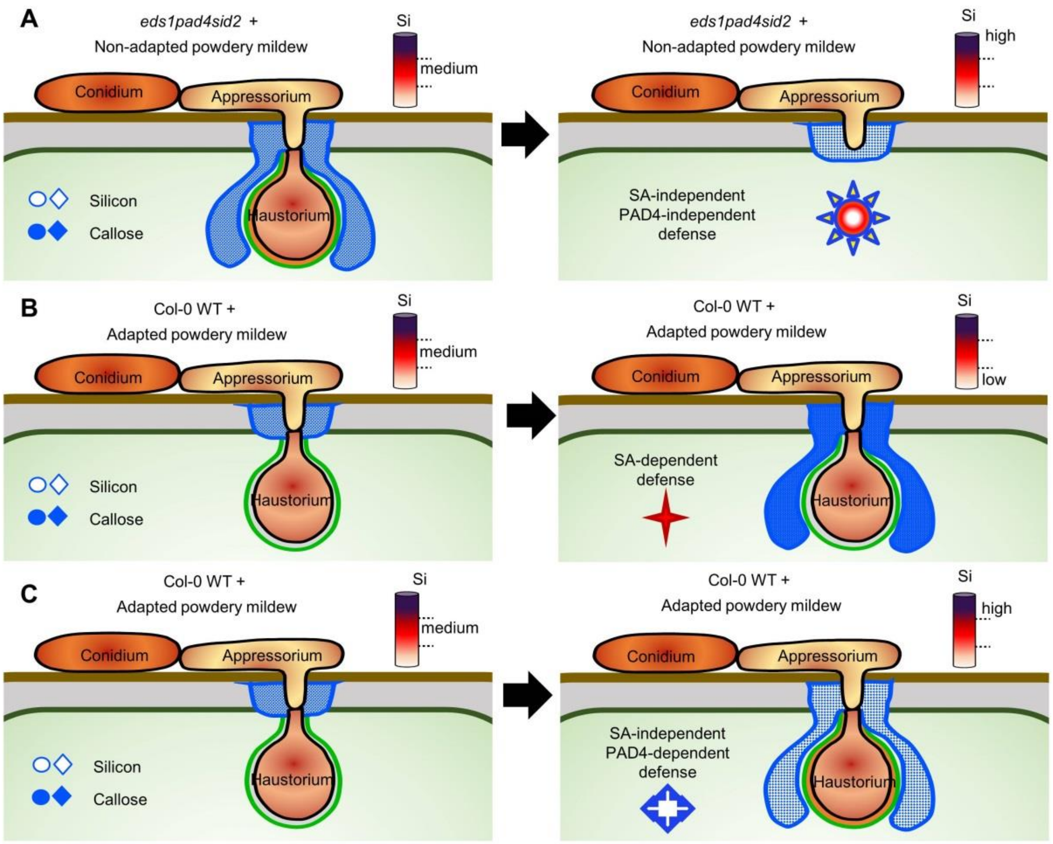


**Figure S9. Schematic diagram describing the three scenarios where very low Silicon (Si) or high Si confers enhanced resistance to powdery mildew, and the potential synergistic action between Si and callose.**

Based on publish results and our observations from this study, we speculate that Si may polymerize with cell wall components especially callose to fortify the cell wall as a major preformed physical barrier to defend plant cells against cell wall-penetrating non- and poorly-adapted pests such as PM. Compromised resistance against such pathogens due to loss of EDS1, PAD4 and SID2 can be compensated by more Si deposition (**A**). Adapted PM has largely overcome this physical barrier; however, depriving Si or callose from such structures may generate a warning signal to activate SA-dependent defense that can promote the callose-rich papilla to further "grow" to (half)encase the haustorium (Meyer *et al.*, 2009), limiting the growth of the adapted PM (**B**). High Si can strengthen the papilla and the callosic encasement, which requires a PAD4-dependent (but SA-independent) mechanism, resulting in enhanced basal resistance (**C**).

**Table S1** Primers used in this work.

| Gene symbol | Primer name | Sequence 5'–3' |
|---|---|---|
| | 35S-F | TCCTTCGCAAGACCCTTCCTCTAT |
| LOC103487002 | CmeLsi1-F | caccATGAGTTCTATAAATCCTGAGCT |
| LOC103487002 | CmeLsi1-R | TCAGTGATTGCTTTCGCTAAC |
| LOC100301576 | HvLsi1-F | caccATGGCCAGCAACTCGAGAT |
| LOC100301576 | HvLsi1-R | TCAGACGGGGATGTGGTC |
| AT4G27960 | AtUBC9-F | CAGTGGAGTCCTGCTCTCACAA |
| AT4G27960 | AtUBC9-R | CATCTGGGTTTGGATCCGTTA |
| AT2G14610 | AtPR1-F | AGAGGCAACTGCAGACTCATACAC |
| AT2G14610 | AtPR1-R | AGCCTTCTCGCTAACCCACAT |
| AT5G44420 | AtPDF1.2-F | TGTTCTCTTTGCTGCTTTCGACGC |
| AT5G44420 | AtPDF1.2-R | TGTGTGCTGGGAAGACATAGTTGC |
| AT4G03550 | AtPMR4-F | ATCTTTACCCGTGGTGATGC |
| AT4G03550 | AtPMR4-R | TATGCTCCCTGACACCAAGA |
| AT3G52430 | AtPAD4-F | TTATCCTCCGATGAACCTCTACC |
| AT3G52430 | AtPAD4-R | AAGCCAAAGTGCGGTGAAAG |
